# Loss of Cohesin regulator PDS5A reveals repressive role of Polycomb loops

**DOI:** 10.1101/2021.12.15.472841

**Authors:** Daniel Bsteh, Hagar F. Moussa, Georg Michlits, Ramesh Yelagandula, Jingkui Wang, Ulrich Elling, Oliver Bell

## Abstract

Polycomb Repressive Complexes 1 and 2 (PRC1, PRC2) are conserved epigenetic regulators that promote transcriptional silencing. PRC1 and PRC2 converge on shared targets, catalyzing repressive histone modifications. In addition, a subset of PRC1/PRC2 targets engage in long-range interactions whose functions in gene silencing are poorly understood. Using a CRISPR screen in mouse embryonic stem cells, we identified that the cohesin regulator PDS5A links transcriptional silencing by Polycomb and 3D genome organization. PDS5A deletion impairs cohesin unloading and results in derepression of subset of endogenous PRC1/PRC2 target genes. Importantly, derepression is not associated with loss of repressive Polycomb chromatin modifications. Instead, loss of PDS5A leads to aberrant cohesin activity, ectopic insulation sites and specific reduction of ultra-long Polycomb loops. We infer that these loops are important for robust silencing at a subset of Polycomb target genes and that maintenance of cohesin-dependent genome architecture is critical for Polycomb regulation.

## INTRODUCTION

In metazoans, precise epigenetic regulation of gene expression enables the development of diverse cell types despite the same underlying genomic blueprint. Gene expression is primarily controlled by DNA-binding transcription factors directing the transcriptional apparatus. However, epigenetic mechanisms modulate chromatin to directly and indirectly regulate transcription. Polycomb repressive complexes are chromatin-modifiers and serve as prototypes of epigenetic gene regulation via histone modifications. Decades of research have cemented their roles in establishing and maintaining cell identity throughout development across organisms, ranging from *Drosophila melanogaster* to vertebrae ^1, 2^. Moreover, aberrant activity of Polycomb complexes and other epigenetic regulators contribute to diverse diseases, including cancer initiation and metastasis, highlighting the importance of understanding the pathological mechanisms ^3–10^.

Polycomb group (PcG) proteins are conventionally grouped into Polycomb repressive complexes 1 and 2 (PRC1 and PRC2). All PRC1 complexes share RING1A/B as their catalytic core subunit, which deposits H2AK119ub at its targets, whereas PRC2 contains the histone methyltransferase EZH1/2, which catalyzes H3K27me3 at targets ^11–14^. In vertebrates, PRC1 complexes have diversified into distinct subcomplexes based on incorporation of one of six paralogous PCGF proteins (PCGF1-6). PCGF2 or PCGF4 dictate assembly of canonical PRC1 (cPRC1) which specifically incorporates CBX (chromobox-containing protein) subunits. CBX subunits endow cPRC1 with the capacity to bind H3K27me3, which promotes cPRC1 recruitment to PRC2 target genes and transcriptional silencing ^11, 15, 16^. PCGF1, 3, 5 and 6 form variant PRC1 (vPRC1) complexes which harbor RING1 and YY1-binding protein (RYBP), or its paralogue YAF2 instead of CBX, and rely on PRC2-independent mechanisms of chromatin targeting. Thus, PRC1 and PRC2 are generally considered to exert their repressive functions in a synergistic manner, but they have different mechanisms of targeting, signaling and repression (reviewed in ^17^).

Recent studies have established that vPRC1 can act upstream of PRC2 and cPRC1, and that its H2AK119ub deposition is critical for Polycomb-dependent gene silencing ^18–20^. Indeed, loss of PRC2 or cPRC1 does not substantially compromise the repression of Polycomb target genes in mouse embryonic stem cells (mESCs) expressing vPRC1 ^18, 19^. These findings have propelled vPRC1 to the center of coordinating and establishing a repressive Polycomb chromatin domain. Although cPRC1 contributes minimally to H2AK119ub deposition, it possesses the unique capacity to mediate long-range 3D interactions between Polycomb target genes ^12, 21–27^, which has been shown to contribute to gene silencing in flies ^28^.

The redundant functions of vPRC1 and cPRC1 complicate dissecting their individual mechanisms by genetic analysis ^18, 19, 29, 30^. To circumvent this limitation, we previously developed a Polycomb *in vivo* Assay that reports the activity of distinct PRC1 complexes. Briefly, we generated mESCs that can recruit ectopic cPRC1 or vPRC1 to an integrated TetO repeat flanked by fluorescent reporters (Fig. 1a) ^31^. For instance, ectopic expression of a CBX7-Tet repressor domain (TetR-CBX7) fusion triggers the assembly of cPRC1 at the TetO sites, Polycomb-dependent histone modifications and reporter gene silencing. Binding of the TetR fusion is released upon addition of Doxycycline (Dox), and we found that more than 70% of cells maintained cPRC1-induced, but not vPRC1-induced, silencing in the presence of Dox. We showed that sequence-independent propagation of cPRC1-induced silencing requires H3K27me3 and H2AK119ub, suggesting that it relies on PRC1/PRC2 feedback. However, the mechanism and players required for heritable PRC1/PRC2-mediated silencing, and whether they are the same at all target genes, remain incompletely understood.

**Fig. 1.**
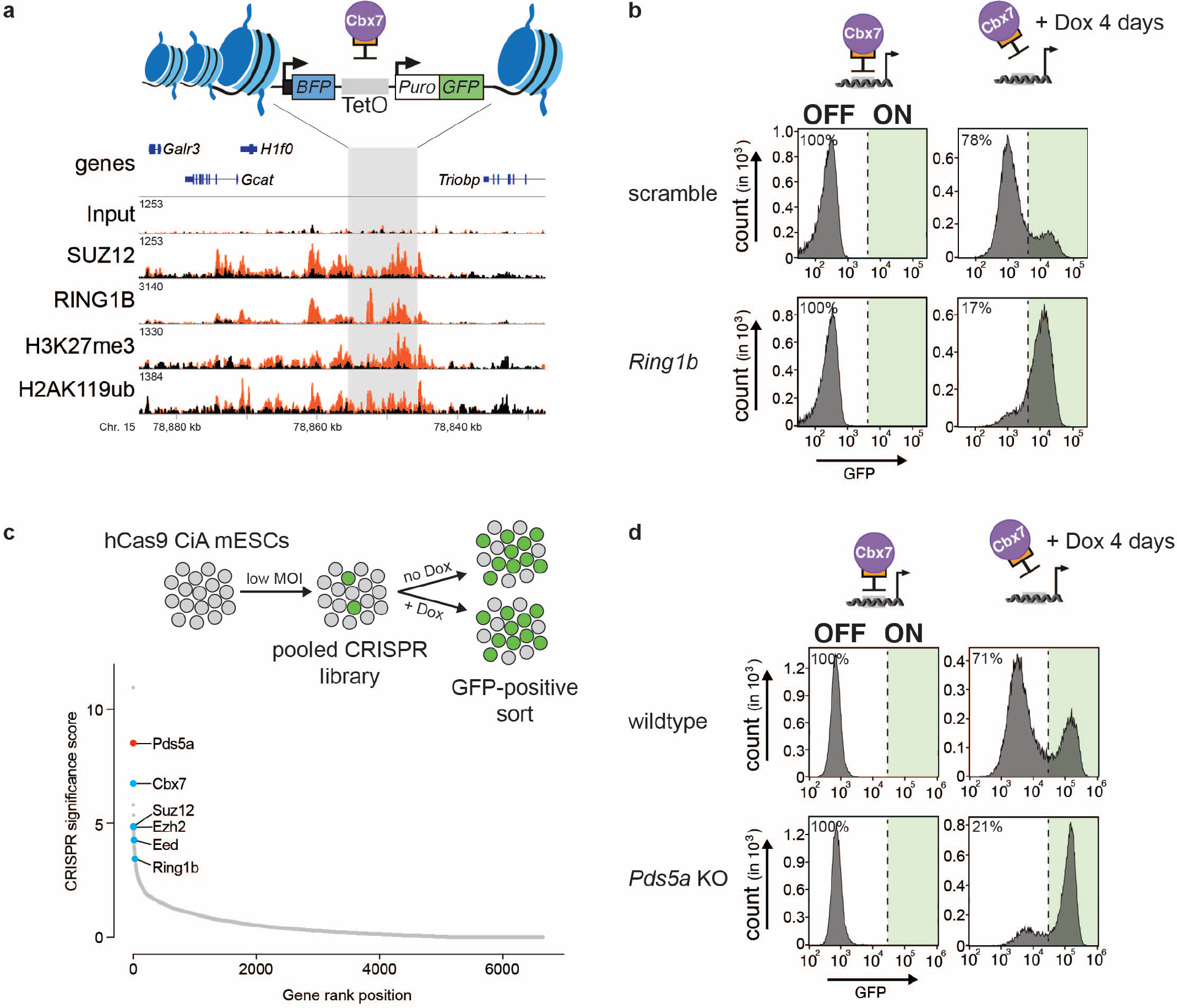
CRISPR screen of cPRC1-dependent gene silencing reveals PDS5A. **a,** Schematic of ectopic dual reporter locus consisting of 7x TetO landing sites flanked by an upstream Ef1a promoter driven BFP and a downstream PGK driven puromycin/GFP. Underneath shown are genomic ChIP-CapSeq screenshot of polycomb proteins and histone modifications before (black) and after TetR-Cbx7 expression (orange). **b,** Flow cytometry histograms of GFP reporter expression activity at continuous binding of TetR-Cbx7 fusion proteins (left) and after doxycyline induced release from the locus for 6 days (right). Top panel shows propagation of silencing after CRISPR-Cas9 infection utilizing a scrambled control sgRNA, and bottom panel CRISPR-Cas9 with sgRNA directed against Ring1b. Percentages refer to fraction of GFP-negative cells. **c,** Schematic of CRISPR screen design and flow cytometry sorting scheme. MOI = multiplicity of infection. Below, all nuclear factors genes targeted in the CRISPR screen (sgRNA library described in ^32^) are ranked by their corresponding CRISPR significance score (-log 10 MAGeCK significance score ^32^; n = mean of three independent experiments)). **d,** Flow cytometry histograms of GFP reporter expression before and after doxycycline-induced release of TetR-Cbx7 binding for 4 days in wildtype reporter cells and FACS sorted Pds5a-sgRNA CRISPR-Cas9 targeted population. Percentages refer to fraction of GFP-negative cells.

Here we performed a CRISPR-mutagenesis screen to identify novel regulators of silencing by PRC1/PRC2. We discovered that the cohesin regulator PDS5A is required for repression of canonical Polycomb target genes. Unexpectedly, loss of PDS5A does not substantially impact repressive Polycomb chromatin domains, but instead disrupts ultra-long chromatin loops between Polycomb target genes. Our work uncovers a subset of Polycomb target genes that require distal 3D interactions for transcriptional silencing.

## RESULTS

### CRISPR screen of cPRC1-induced gene silencing reveals Pds5a dependence

To identify novel regulators of PRC1/PRC2-mediated target gene silencing, we performed CRISPR-based screening. As a screening platform, we introduced stable expression of hCas9 into our TetR-CBX7 reporter line. Parental lines established a Polycomb chromatin domain with high levels of RING1B, SUZ12, H3K27me3 and H2K119ub surrounding the TetO nucleation site (Fig. 1a and Extended Data Fig. 1a). As a proof-of-principle, we infected this line with lentiviral vectors expressing either scramble sgRNA or sgRNA specific for *Ring1b*. We found that CRISPR mutation of *Ring1b* had a negligible effect on silencing induced by TetR-CBX7 (in the absence of Dox), but strongly impaired the epigenetic maintenance of silencing (in the presence of Dox, Fig. 1b), consistent with our previous observations ^31^. Thus, our TetR-CBX7 reporter mESCs recapitulate epigenetic Polycomb-dependent gene silencing and are sensitive to genetic perturbations.

Using this platform, we performed pooled CRISPR screens with unique molecular identifiers (UMIs), which allow analysis of mutant phenotypes at a single-cell level ^32^. The UMI CRISPR library contained approx. 27,000 sgRNAs targeting all annotated mouse nuclear protein-coding genes with four sgRNAs per gene. Each sgRNA was paired with thousands of barcodes representing UMIs, improving the signal-to-noise ratio and hit calling. hCas9-expressing TetR-CBX7 reporter mESCs were transduced with the pooled library and selected with neomycin. We used FACS to isolate GFP-positive cells (Fig. 1c and Extended Data Fig. 1b), and the unsorted population served as background control. Because GFP activation occurs at a very low frequency, we performed repeated FACS in the screen to enrich for GFP-positive cells. Relative enrichment of sgRNAs was determined by sequencing of UMIs in both populations followed by statistical analysis using MAGeCK ^33^.

We performed a screen with reporter cells cultured without Dox (Fig. 1c), and uncovered 51 genes that were significantly enriched in the GFP-positive cell population (p-value < 0.005) (Supplementary Table 1). We also performed a separate screen of cells treated with Dox for 3 days, but spontaneous GFP re-activation in some cells, independently of any mutation, precluded us from identifying statistically significant hits. The top hits in our screen included genes that encode subunits of cPRC1 (*Cbx7, Ring1b*) and of PRC2 (*Ezh2*, *Suz12*, *Eed*), indicating that our screening approach identified known genes required for TetR-CBX7-induced silencing (*Cbx7*) and for the epigenetic maintenance of silencing (*Ring1b, Suz12*) (Fig. 1c). Notably, *Pds5a*, which encodes a regulator of the cohesin complex, was the second most-significant hit in the screen. To validate this hit, we used CRISPR-Cas9 to target *Pds5a* independently, and observed reduced silencing in TetR-CBX7 reporter mESCs treated with Dox (Fig. 1d and Extended Data Fig. 1c). Thus, similar to *Ring1b*, *Pds5a* is required for the epigenetic maintenance of silencing induced by cPRC1.

The cohesin protein complex is composed of three core subunits, SMC1, SMC3 and RAD21 (also known as SCC1), which form a tripartite ring structure that entraps DNA^34^. Several auxiliary cohesin proteins are critical for dynamic regulation of DNA interactions. For instance, cohesin release from the DNA involves STAG1/2, WAPL and PDS5A/B which associate at the interface between SMC3 and RAD21 and control ring opening ^35–43^. In addition to *Pds5a*, our CRISPR screen without Dox revealed enrichment of *Stag2*, albeit below the significance cutoff (p-value = 0.045). Notably, a recent study linked STAG2 to Polycomb domain compaction ^44^, further supporting a potential role of cohesin regulation in Polycomb-dependent gene silencing.

Overall, our CRISPR screen suggests that the regulation of genome topology and cohesin by *Pds5a* promotes Polycomb-induced silencing.

### Loss of Pds5a results in de-repression of endogenous PRC1/PRC2 target genes

To determine the impact of PDS5A deletion on endogenous Polycomb-dependent gene regulation, we generated *Pds5a* knockout mESCs using CRISPR-Cas9 (*Pds5a* KO). In addition, we obtained a loss-of-function (LOF) mESC line harboring a disruptive gene-trap in the second intron of the *Pds5a* gene (*Pds5a*^GT^ KO) ^45^. Since gene-trap disruption is reversible, we also generated a matched control mESC line in which *Pds5a* expression was restored (*Pds5a*^GT^ WT). *Pds5a* knockouts as well as the rescue were confirmed by western blot (Fig. 2a). PDS5A deletion did not impact the abundance of the cohesin subunit SMC3, of the PcG proteins SUZ12 and RING1B, nor the global levels of their associated histone modifications (Fig. 2a).

**Fig. 2.**
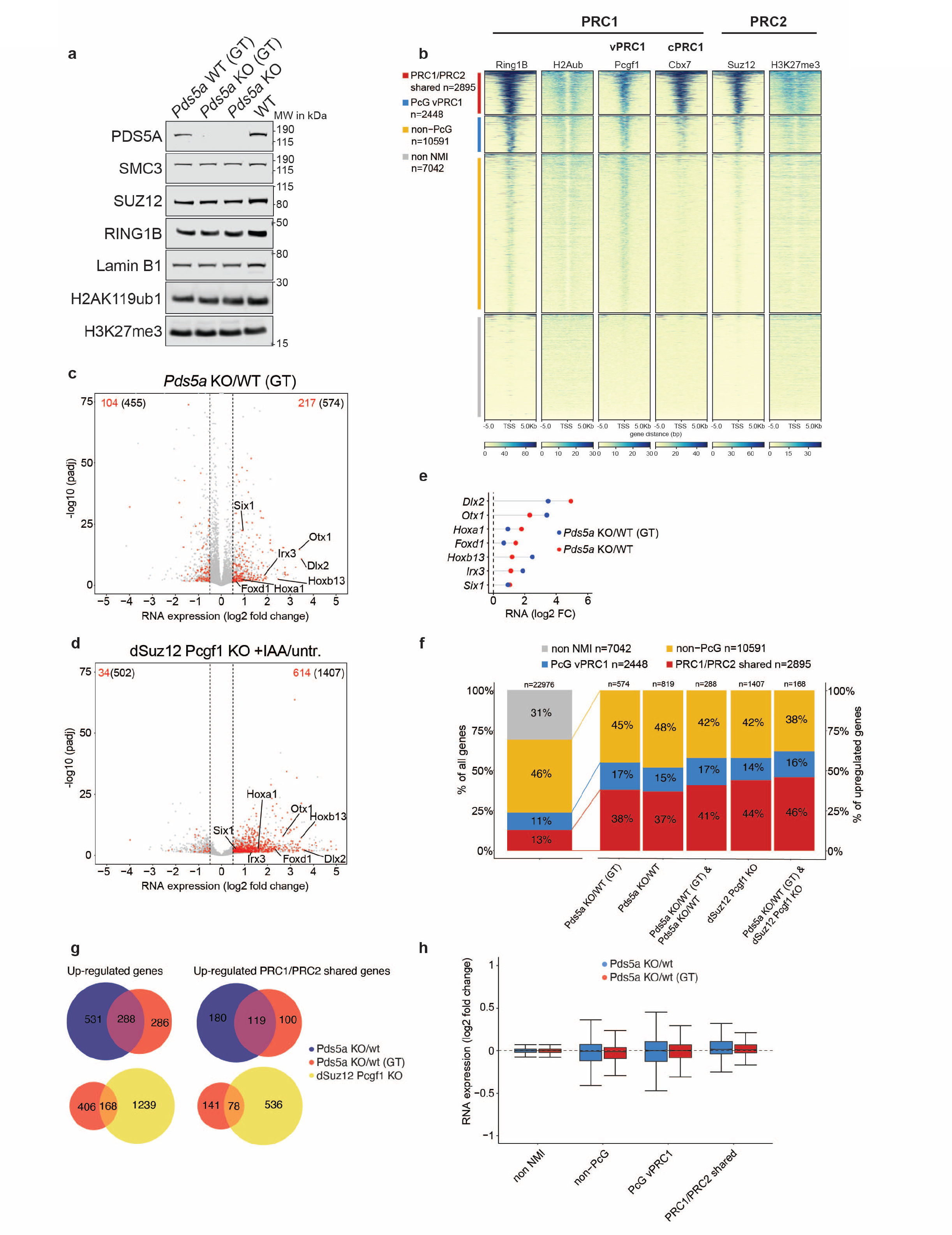
Loss of cohesin regulator PDS5A results in de-repression of canonical PRC1/PRC2 target genes. **a,** Western blot characterization of cohesin and polycomb proteins and histone modifcations in two independent Pds5a KO mESC cell lines and their corresponding wild types. **b,** Calibrated ChIP-seq heatmaps of PRC1 proteins (Ring1B, Pcgf1 (variant PRC1), Cbx7 (canonical PRC1)) and its histone modification H2AK119ub, as well as PRC2 proteins and histone modification Suz12 and H3K27me3, respectively. cChiP-seq signal is plotted around the TSS (+/-5kb). Genes classified as PRC1/PRC2 shared genes (red; n=2895) yield a Bio-Cap peak (Data from ^55^) for non-methylated CpG island, Ring1b peak for PRC1 and a Suz12 peak for PRC2 within 3kb of the TSS. PcG vPRC1 classified genes (blue; n=2448) have only a Ring1b peak within 3kb of the TSS, and genes with no Ring1b or Suz12 peak are fall in the class of non-PcG (yellow;n=10591). Genes without a non-methylated CpG island (NMI) are in class non-NMI (n = 7042). **c,** Volcano plot of RNA-seq expression change in Pds5a genetrap (GT) KO vs. wild type mESCs. Genes from the PRC1/PRC2 shared class (n=2895) with increased or decreased expression are marked in red (padj < 0.05 and log2 fold change > 0.5 or < -0.5). Total numbers of up-/and down-regulated genes are indicated in the top left and right corners (PRC1/PRC2 shared class in red and of all genes in black). **d,** Volcano plot of RNA-seq expression change in 48 hrs IAA-treated vs. untreated dSuz12 Pcgf1 KO mESCs. Genes from the PRC1/PRC2 shared class (n=2895) with increased or decreased expression are marked in red (padj < 0.05 and log2 fold change > 0.5 or < - 0.5). Total numbers of up-/and down-regulated genes are indicated in the top left and right corners (PRC1/PRC2 shared class in red and of all genes in black). **e,** Dotplots of log2 fold changes of selected representative genes from the PRC1/PRC2 shared genes class in both independent Pds5a KO mESC cell lines. **f,** Stacked bargraphs of percent gene distribution between classes of all genes (left) and of up-regulated genes in both Pds5a KO cell lines, dSuz12 Pcgf1 KO IAA treated cells and in the overlap of Pds5a KO (GT) and dSuz12 Pcgf1 KO. **g,** Euler diagrams of overlap of all, as well as only PRC1/PRC2 shared up-regulated genes between both Pds5a KO cell lines (top) and between Pds5a KO (GT) and dSuz12 Pcgf1 KO. **(H)** Boxplots of RNA-seq log2 fold change expression changes for each gene class for both Pds5a KO mESC cell lines.

Given cohesin’s essential role in sister chromatid cohesion, deletion of cohesin subunits frequently impairs cell proliferation, hampering the analysis of its precise function in gene regulation ^46–48^. Although most cohesin proteins are essential for mESC viability ^46–48^, both PDS5A and STAG2 have paralogs with redundant but not identical functions that are sufficient to maintain self-renewal and proliferation ^35^. Indeed, *Pds5a* KO mESC lines displayed characteristically dense colonies that could be stably maintained in culture, similar to wildtype mESCs (Extended Data Fig. 2a). Consistently, cell cycle profiles and pluripotency marker expression were highly comparable between KO and control mESCs, suggesting that PDS5A is largely dispensable for mESC self-renewal and proliferation (Extended Data Fig. 2a-c).

To evaluate how endogenous Polycomb target genes are affected by PDS5A deletion, we first categorized all transcription start sites (TSSs) in mESCs based on the occupancies of PRC1 and PRC2, and the enrichment of their associated histone modifications. In addition, because non-methylated CpG-islands (NMIs) have emerged as a general feature of Polycomb target genes ^49–54^, we used available BioCap data to distinguish NMI TSSs from methylated, CpG-poor TSSs which we annotated “non-NMI”^55^. We classified NMI TSSs with overlapping peaks of RING1B and SUZ12 as “shared PRC1/PRC2 target genes”, whereas NMI TSSs with RING1B peaks only were classified as “vPRC1 target genes”. Notably, “shared PRC1/PRC2 target genes” displayed high levels of H3K27me3 and were predominantly bound by CBX7-containing cPRC1. In comparison, CBX7 was low at vPRC1 target genes which showed substantial PCGF1 occupancy instead. NMI TSSs lacking both RING1B (PRC1) and/or SUZ12 (PRC2) peaks within 3 kb of their TSSs were classified as “non-PcG target genes” (Fig. 2b).

Transcriptome profiling of *Pds5a*^GT^ KO mESCs and *Pds5a* KO mESCs revealed differential expression of ∼1000 and ∼1500 genes, respectively (cutoff: LFC +/-0.5, padj. 0.05) (Fig. 2c, g, h and Extended Data Fig. 2d, g). Notably, Gene Ontology terms of upregulated genes were related to developmental processes such as neurogenesis and pattern specification, reminiscent of Polycomb target genes (Extended Data Fig. 2e and Extended Data Fig. 2f). Although only 13% of expressed genes in wildtype mESCs were shared PRC1/PRC2 target genes, this group represented 38% and 37% of the upregulated genes in *Pds5a*^GT^ KO and *Pds5a* KO mESCs, respectively (Fig. 2c). We infer that silencing of this class is particularly dependent on PDS5A function. We did not observe a bias in the relative distribution of gene classes among downregulated genes, suggesting that PDS5A specifically promotes repression of shared PRC1/PRC2 target genes (Extended Data Fig. 2h, i).

To examine the role of PDS5A versus PRC1/PRC2 in the regulation of PRC1/PRC2 target gene expression, we took advantage of an mESC line that lacks the vPRC1 component PCGF1 and offers inducible degradation of the PRC2 component SUZ12 (*dSuz12^Pcgf^*^1^*^-null^*). PRC2, cPRC1 and vPRC1, which act redundantly, are all impaired upon auxin treatment of *dSuz12^Pcgf^*^1^*^-null^* mESCs. Transcriptome analysis of *dSuz12 ^Pcgf^*^1^*^-null^* mESCs treated with IAA for 48h revealed significant changes in expression of ∼1900 genes (cutoff: LFC +/-0.5, padj. 0.05). More than two-thirds of dysregulated genes (∼1400) displayed increased expression, consistent with loss of PRC1/PRC2-mediated gene silencing in mESCs. Importantly, similar to *Pds5a* KO mESCs, shared PRC1/PRC2 target genes accounted for 44% of upregulated genes. Preferential enrichment of this class was specific, as the fractions of vPRC1- and non-PcG target genes remained unchanged (Fig. 2d, f, g).

These results reveal that PDS5A plays an important role in transcriptional silencing of endogenous, shared PRC1/PRC2 target genes, validating our CRISPR screening results.

### PDS5A deletion has minimal effect on Polycomb repressive chromatin domains

Polycomb-mediated transcriptional silencing is linked to the formation of repressive chromatin domains marked by PRC1 and PRC2 occupancy and deposition of Polycomb-dependent histone modifications ^2, 17^. We considered that impaired silencing of endogenous PRC1/PRC2 target genes upon PDS5A deletion results from erosion of these repressive chromatin domains. To test this hypothesis, we used calibrated ChIP-seq (cChIP-seq) to compare enrichment of RING1B, SUZ12, H3K27me3 and H2AK119ub between wild type and Pds5a^GT^ KO mESCs. PDS5A deletion led to a modest reduction in RING1B and H3K27me3 signals at shared PRC1/PRC2 and vPRC1 target genes, but no differences in SUZ12 and H2AK119ub enrichment across the different gene classes upon (Fig. 3 a, c, e, f and Extended Data Fig. 3a, c, f). In addition, we used ATAC-seq analysis to analyze wildtype and Pds5a^GT^ KO mESCs but did not observe significant differences in DNA accessibility, arguing that the modest difference in histone modifications does not affect the integrity of Polycomb repressive chromatin domains (Fig. 3g). Given the substantial derepression of shared PRC1/PRC2 target genes, the limited effects on repressive chromatin domains are surprising and in stark contrast with the dramatic reduction of RING1B and H2AK119ub in IAA-treated *dSuz12 ^Pcgf^*^1^*^-null^*, which have similar silencing defects as PDS5A depletion (Fig. 3b, d and Extended Data Fig. 3b, d, e).

**Fig. 3.**
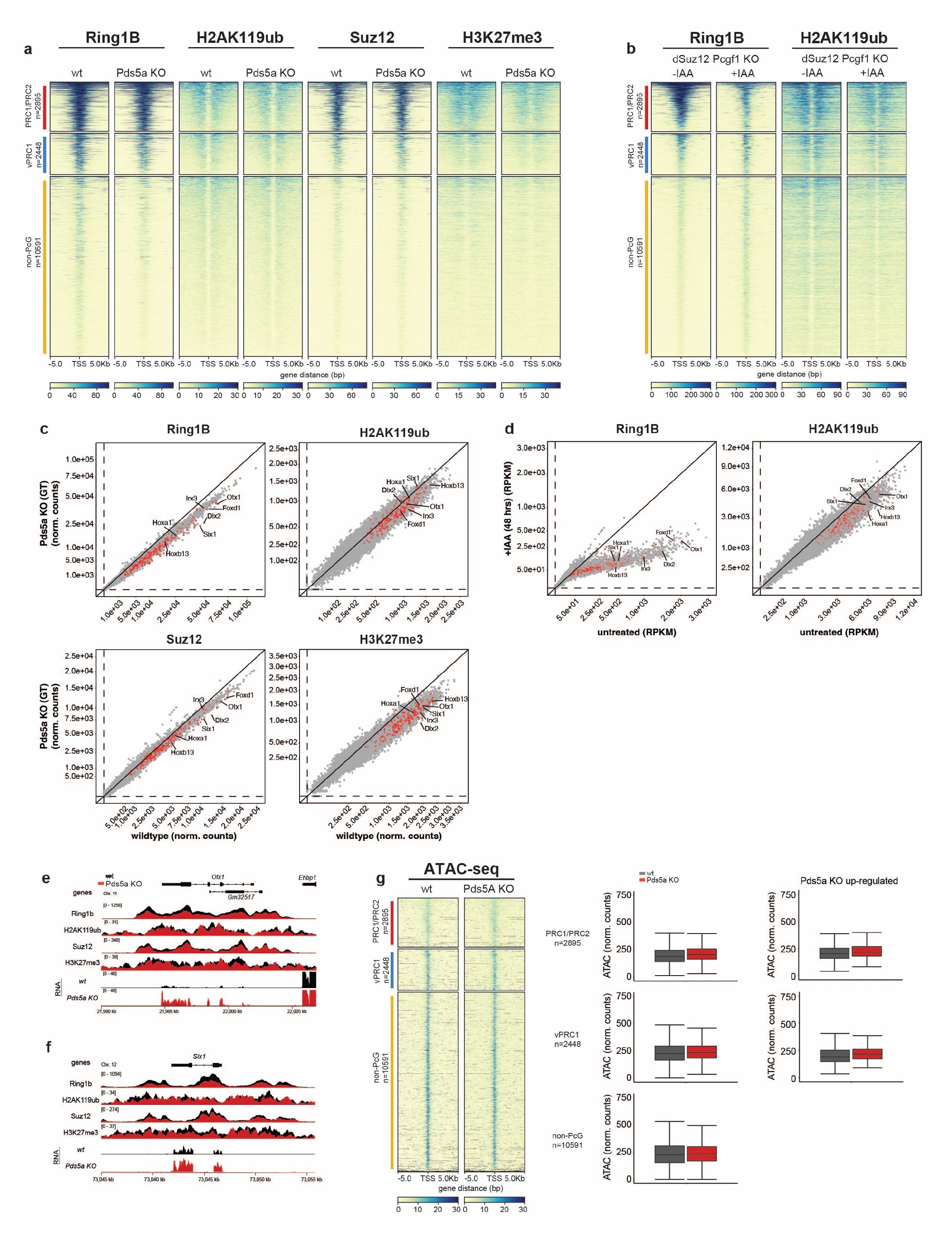
PDS5A deletion has minimal effect on Polycomb repressive chromatin domains. **a,** Wildtype and Pds5a KO cChIP-seq heatmaps of PRC1 (Ring1b and H2AK119ub) and PRC2 (Suz12 and H3K27me3). cChiP-seq signal is plotted around the TSS (+/- 5kb). Heatmap is divided in 3 gene classes (see Fig2b): PRC1/PRC2 shared genes (red; n=2895), PcG vPRC1 (blue; n=2448), non-PcG (yellow; n=10591). **b,** cChIP-seq heatmaps of PRC1 (Ring1b and H2AK119ub) in 48 hrs IAA-treated vs. untreated dSuz12 Pcgf1 KO mESCs. **c,** Wild type and Pds5a KO scatterplots of cChIP signal around TSS for each gene (Sum of TSS +/- 2.5 kb). Genes from the PRC1/PRC2 shared gene class, which are up-regulated in Pds5a KO cells (see Fig 2c) are marked in red and selected representative genes are labelled individually (see Fig 2d). **d,** cChIP scatterplots as in (c) for 48 hrs IAA-treated vs. untreated dSuz12 Pcgf1 KO mESCs. **e,** IGV genomic cChIP-seq track screenshot of RNA-seq, PRC1 and PRC2 polycomb proteins and histone modifications in wildtype (black) and Pds5a KO (red) mESCs at the *Otx1* locus and **f,** at the *Six1* locus. **g,** ATAC-seq heatmap, boxplots for each gene class (see Fig. 3a) and the corresponding subset of transcriptionally up-regulated genes.

Together, these results show that PDS5A depletion has limited impact on the formation of repressive chromatin domains. The minimal reduction of PcG protein binding and histone modifications appears disproportionate to the degree of aberrant transcriptional activation at shared PRC1/PRC2 target genes upon loss of PDS5A, suggesting that *Pds5a* KO impairs other mechanisms that are critical for transcriptional silencing.

### PDS5A colocalizes with cohesin and destabilizes chromatin binding

The PDS5A/B interaction with WAPL is critical for transient opening of the cohesin ring to release DNA loops during interphase ^35–43^. We used cChIP-seq to determine the genomic distribution of PDS5A relative to the cohesin subunit RAD21, CTCF and PcG proteins in wildtype mESCs. As expected, PDS5A showed extensive overlap with RAD21 and CTCF (Extended Data Fig. 4d-e and Extended Data Fig. 4g) ^56, 57^. In contrast, PDS5A was absent from Polycomb target genes, suggesting that PDS5 indirectly promotes silencing of shared PRC1/PRC2 target genes.

**Fig. 4.**
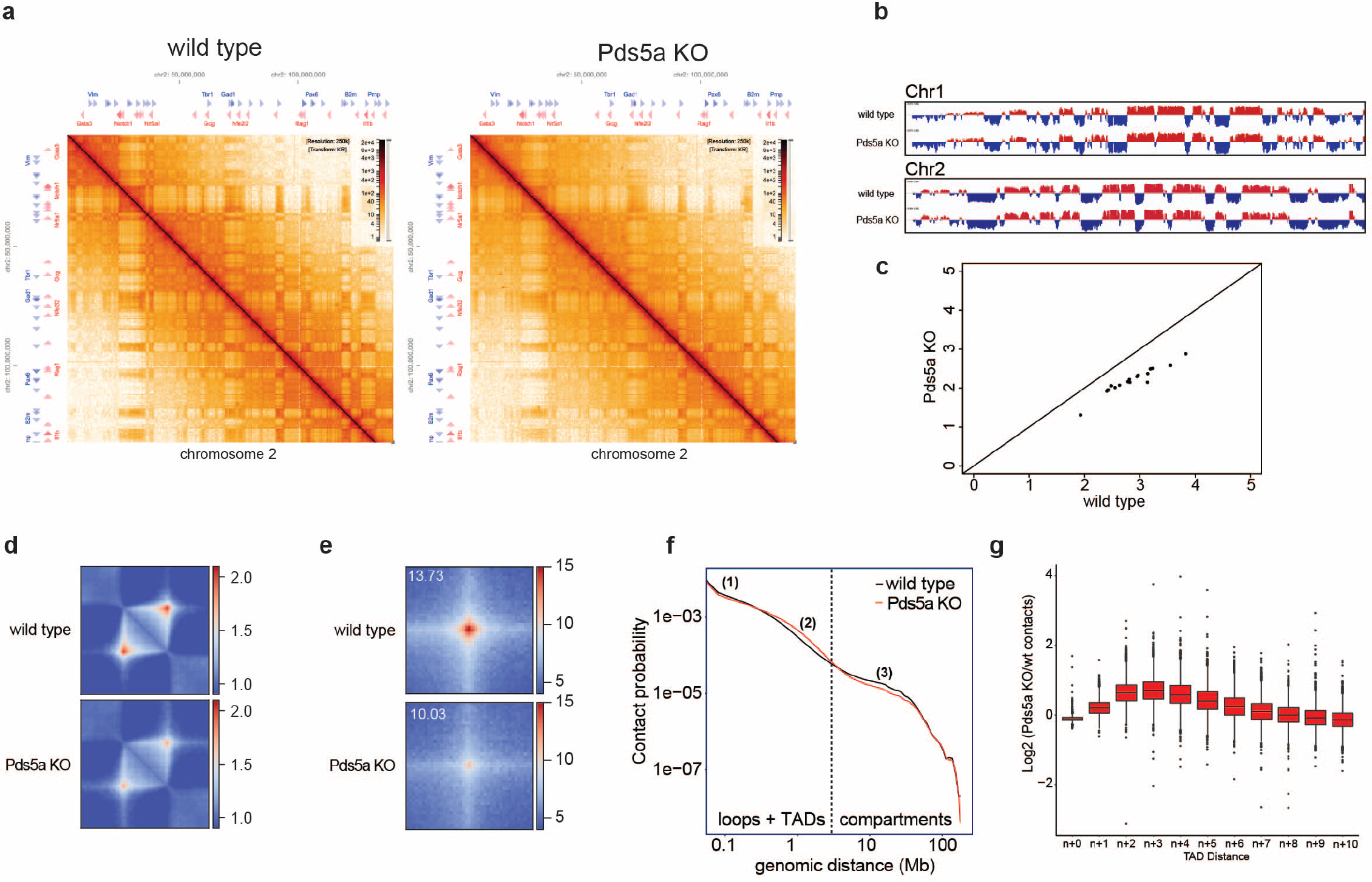
PDS5A deletion increases cohesin-dependent loop extrusion causing TAD boundary violations. **a,** KR normalized Hi-C contact matrices of chromosome 2 in wild type (left) and Pds5a KO (right) mESCs. **b,** Eigenvector compartment signal tracks at 250 kb bin resolution are plotted for both wild type and PDS5A. **c,** Wild type vs. Pds5a KO compartment strength (generated in GENOVA) are plotted. One dot for each chromosome. **d,** Aggregate Hi-C pileup analysis (observed/expected enrichment) using coolpup.py from wild type (top panel) and Pds5a KO (bottom panel) at wild type mESC TADs called in Bonev et al. 2017 ^59^ mESC dataset (5kb resolution). **e,** Aggregate Hi-C pileup analysis (observed/expected enrichment) using coolpup.py from wild type (left panels) and Pds5a KO (right panels) at wild type mESC loops +/- 100kb called in Bonev et al. 2017 ^59^ mESC dataset (5kb resolution). Bottom panels show analysis for subset of only Cohesin dependent (CTCF anchored) and PRC1 dependent (Ring1b anchored) loops. In each pileup plot, the mean enrichment scores of the three center pixels is displayed in the top left corner. **f,** RCP plot (Relative contact probability) depicting genomic distance dependent contact frequency for wild type and Pds5a KO mESCs. Dashed line indicates expected size range of TADs and compartments. (1) Relative reduction of short-range contacts (0-500kb) in Pds5a KO; (2) Relative increase of mid-range contacts (500kb-5mb) in Pds5a KO; (3) Relative reduction of long-range contacts (>5mb) in Pds5a KO. **g,** Quantification of TAD border violation. Boxplot of log2 change in HiC contacts with n+ (x-axis) neighboring TADs between wild type and Pds5a KO.

Depletion of WAPL or of both PDS5A and PDS5B, increases the cohesin residence time on chromatin, resulting in continued loop extrusion, formation of larger TADs and loss of compartmentalization ^37, 42^. To evaluate potential changes in the genomic enrichment of cohesin between wildtype and Pds5a^GT^ KO mESCs, we performed RAD21 cChIP-seq (Extended Data Fig. 4d-g). Peak calling revealed three groups of RAD21 binding sites: “common” sites in wildtype and Pds5a^GT^ KO mESCs (n=15941), sites that scored in “wildtype only” (n=4288), and sites that scored in “Pds5a^GT^ KO only” (n=6789). Importantly, “common” binding sites showed increased RAD21 occupancy in Pds5a^GT^ KO mESCs compared to wildtype, consistent with previous observations in HeLa cells (Wutz et al., 2017).

Together, these results suggest that PDS5A acts at CTCF binding sites to release cohesin from chromatin in mESCs.

### PDS5A deletion increases cohesin-dependent loop extrusion, causing TAD boundary violations

Based on our findings above, we hypothesized that loss of PDS5A leads to a cohesin-dependent dysregulation of 3D genome architecture that compromises the silencing of endogenous, shared PRC1/PRC2 shared target genes. To investigate if PDS5A deletion alters the 3D genome architecture, we performed *in-situ* Hi-C on wildtype and *Pds5a* KO mESCs. After quality control, we combined sequencing reads of Hi-C replicates amounting to a total of ∼365 million valid unique *cis*-contacts per genotype (Supplementary Table 2). We observed decreased interaction frequencies in Knight-Ruiz (KR) ^58^ normalized Hi-C contact matrices resulting in reduced “checkerboard” patterns of alternating A and B compartments (Fig. 4a). Based on eigenvector analysis, compartment signal was reduced, but compartment did not switch from A to B or vice versa (Fig. 4b and Extended Data Fig. 4a, b). Reduced compartmentalization in *Pds5a* KO mESCs was further confirmed by their lower compartment strength, a related benchmark measuring interactions within compartments (A/A or B/B) compared to between compartments (A/B or B/A) (Fig. 4c). When examining relative contact probabilities (RCP) as a function of genomic distance, we found reduced compartmentalization, manifested by a decrease in very long long-range contacts (>5 Mb) in *Pds5a* KO mESCs relative to wildtype. Contact probabilities in the relative short-range (50-500 kb) were also slightly reduced. In contrast, interactions in the mid- to long-range (500 kb - 5 Mb) were increased in *Pds5a* KO mESCs relative to wildtype (Fig. 4f).

Whereas PDS5A depletion reduced compartmentalization, topologically associating domain (TAD) sizes were on average larger in *Pds5a* KO mESCs compared to wild type mESCs (Extended Data Fig. 4c). To explore differences in TAD structure and loop formation between wildtype and *Pds5a* KO, we utilized publicly available high-resolution mESC Hi-C data to identify high confidence TAD intervals and loops ^59^. Aggregate TAD and loop analysis showed that PDS5A loss resulted in a relative contact reduction at pre-existing wildtype TADs and loops (Fig. 4d, e). We also detected increased interaction frequencies with neighboring TADs in *Pds5a* KO mESCs, whereas intra-TAD interactions were slightly decreased (Fig. 4g). These changes resemble those observed upon WAPL and/or PDS5A/B depletion in cancer cells, where increased cohesin residence time leads to an extension of chromatin loops, resulting in a genome-wide shift towards longer range interactions and violation of TAD boundaries ^37, 42^. Overall, our data suggest that PDS5A loss impairs cohesin unloading in mESCs.

### PDS5A is required to maintain a subset of Polycomb loops

To explore how loss of PDS5A affects long-range interactions between Polycomb target genes, we used Ring1B cChIP-seq to identify 525 Polycomb (PcG) loops in wildtype mESCs (Fig. 5a). This number is similar to 336 persistent interactions identified in cohesin-depleted mESCs ^26^, suggesting that Polycomb-associated long-range interaction account only for a small fraction of the 12425 loops detected in mESCs. Further classification of Polycomb loops based on additional PcG proteins and associated histone modifications revealed that virtually all of them (476/525, 91%) arise from long-range interactions between genomic sites harboring shared PRC1/PRC2 target genes, consistent with recent findings ^26^ (Fig. 5a).

**Fig. 5.**
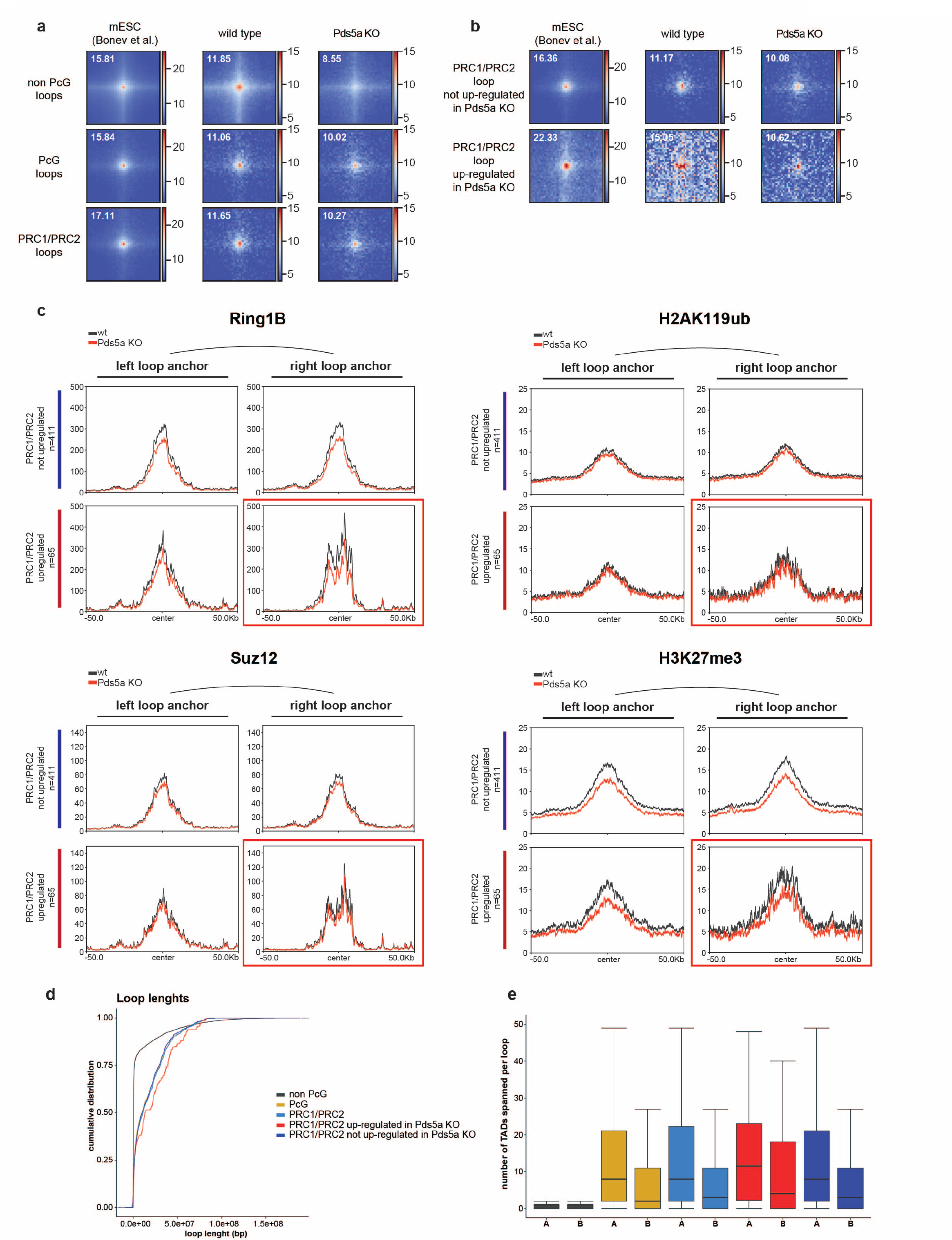
PDS5A is required for maintenance of repressive polycomb loops crossing ultra-long distances. **a,** Loop pileup analysis of HiC data of Bonev et al. 2017 ^59^ mESCs, wildtype and Pds5a KO at not PcG loops, PcG loops (between shared PRC1/2 and/or vPRC1 genes) and shared PRC1/PRC2 PcG loops (between shared PRC1/2 genes). **b,** Loop pileup analysis of HiC data of Bonev et al. 2017 ^59^mESCs, wildtype and Pds5a KO at shared PRC1/PRC2 PcG loops with left and right anchor not-upregulated in Pds5a KO and at shared PRC1/PRC2 PcG loops with left anchor not-upregulated in Pds5a KO and right anchor upregulated in Pds5a KO. **c,** cChIP-seq profile plots of PRC1 (Ring1b and H2AK119ub) and PRC2 (Suz12 and H3K27me3) at Polycomb loops between not-upregulated PRC1/PRC2 shared target genes (left and right loop anchors; n=411) and loops between upregulated PRC1/PRC2 shared target genes (left loop anchor represents the not upregulated side and right loop anchor the Pds5a KO transcriptionally upregulated side; n=65). **d,** Cumulative distribution plots of loop lengths corresponding to loop classes shown in Fig 5A and 5B as well as **e,** number of TADs within the A and B compartment traversed by loops of each class.

Unlike non-Polycomb (non-PcG) loops, which are substantially reduced in *Pds5a* KO mESCs, long-range interactions between shared PRC1/PRC2 target genes displayed on average only minor changes (Fig. 5a). We considered that the aberrant loop extrusion and violation of TAD boundaries that we observed in *Pds5a* KO mESCs could interfere with a subset of Polycomb-associated long-range interactions in a locus-specific manner. Thus, we bifurcated Polycomb loops based on interactions between anchor sites harboring upregulated (65) and not-upregulated (411 – unchanged and downregulated) shared PRC1/PRC2 target genes. Interestingly, anchor sites in wildtype mESCs that involve upregulated shared PRC1/PRC2 target genes are engaged in stronger loops when compared to the anchor sites that do not involve upregulated genes (Fig. 5b). Strikingly, upon PDS5A deletion, the interaction frequency at upregulated anchor sites was dramatically reduced, whereas interactions between not-upregulated anchor sites were relatively unaffected, similar to the class average (Fig. 5b). These results suggest that local dysregulation of cohesin-mediated chromosome architecture interferes with long-range interactions at a subset of shared PRC1/PRC2 target genes, which in turn compromises gene silencing.

### Repressive polycomb loops crossing ultra-long distances are sensitive to cohesin dysfunction

To understand what defines the subset of Polycomb loops that are vulnerable to cohesin dysfunction, we first compared chromatin modifications between anchor sites of upregulated and not-upregulated PRC1/PRC2 target genes. We noticed that at Polycomb loops of upregulated PRC1/PRC2 target genes, only one of the two anchor sites was associated with loss of gene silencing. To investigate potential differences in the repressive chromatin modifications, we separated the two anchor sites into not-upregulated (left) and upregulated (right) and compared PcG protein occupancy and associated histone modifications (Fig. 5c and Extended Data Fig. 5a). Anchor sites of not-upregulated shared PRC1/PRC2 target genes served as the control dataset. Surprisingly, despite differential expression we found that upregulated (right) and not-upregulated anchor sites (left) had comparable repressive chromatin domains with similar reduction in Ring1B occupancy and H3K27me3 upon PDS5A deletion (Fig. 5c and Extended Data Fig. 5a). Chromatin modifications at upregulated anchor sites were also similar to those at anchor sites of not-upregulated shared PRC1/PRC2 target genes. Together, these results corroborate our genome-wide analysis revealing minimal reduction of repressive chromatin modifications at shared PRC1/PRC2 target genes and strongly suggest that reduced long-range interactions and loss of gene silencing in *Pds5a* KO mESCs are largely uncoupled from changes in Polycomb chromatin domains. Next, we explored if sensitivity to cohesin deregulation is linked to the distance between Polycomb anchor sites. Comparison of loop sizes of upregulated and not-upregulated shared PRC1/PRC2 target genes revealed a striking difference: loops between anchor sites of upregulated shared PRC1/PRC2 target genes were substantially longer than loops between anchor sites of not-upregulated or all shared PRC1/PRC2 target genes (Fig. 5d). Not surprisingly, this length bias of loops was also reflected in a greater number of TADs within A and B compartments traversed by anchor sites of upregulated shared PRC1/PRC2 target genes. Based on these results we conclude that ultra-long Polycomb loops are most vulnerable to cohesin dysfunction. We speculate that traversing a greater number of TADs increases the probability of interference by cohesin-mediated loop extrusion and TAD boundary violations. Importantly, by uncoupling loss of Polycomb loops from changes in repressive chromatin modifications, these results reveal a subset of shared PRC1/PRC2 target genes that depend on long-range interactions in silencing.

### Loss of repressive Polycomb loops is linked to cohesin-mediated insulation gain

To uncover the potential mechanism by which increased cohesin residence could interfere with repressive ultra-long loops between shared PRC1/PRC2 target genes, we defined regions in the genome with significant local changes in 3D chromosome architecture. Specifically, we calculated insulation scores ^60^ in 250 kb bins across the genomes of wildtype and *Pds5a* KO mESCs. Insulation scores in most bins (∼41000) were unchanged upon PDS5A deletion (Fig. 6a). Additionally, we identified ∼17000 bins with significant reduction in insulation in *Pds5a* KO mESCs, suggesting loss of TAD boundaries in response to increased cohesin residence time (Extended Data Fig. 6a).

**Fig. 6.**
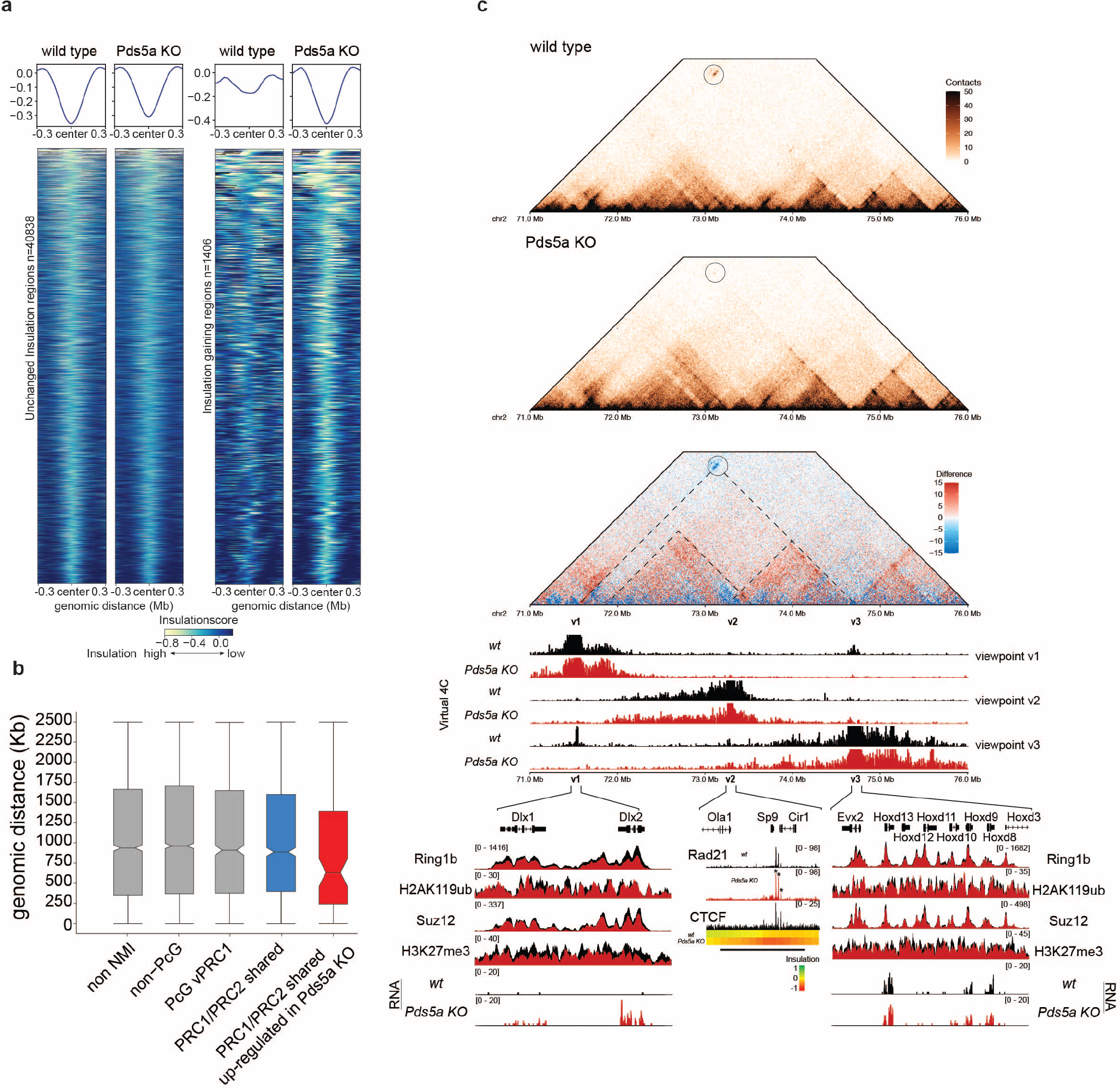
Loss of repressive Polycomb loops is linked to cohesin-mediated insulation gain. **a,** Heatmaps and average profile plots of insulation score of genomic 10kb bins (+/- 300kb) in wild type and Pds5a KO. Left panel shows bins with unchanged insulation (n=40838); Right panel shows insulation gaining bins (n=1408; minimum insulation score decrease of -0.2). **b,** Boxplots for each gene class showing genomic distance (kb) to the closest insulation gaining 10kb bin of the TSS of each gene within its gene class. **c,** Loss of 3D interaction of Dlx1/2 with HoxD cluster. Shown are wild type and Pds5a KO HiC matrices and a differential HiC matrix (blue = loss; red = gain of contacts in Pds5a KO; 10kb resolution and ICE-normalized). Virtual 4C tracks for both wt and Pds5a KO from indicated viewpoints v1-3 are plotted below. Genomic screenshots illustrating polycomb group proteins and histone modification cChIP at Dlx1/2 and the HoxD cluster. Insulation scores, as well as cChIP IGV screenshots of cohesin (Rad21) and CTCF are shown at the Insulation gaining region (indicated by black bar). Change in RNA-seq signal for Dlx2 is plotted underneath.

Intriguingly, we identified ∼1400 genomic regions that gained insulation in *Pds5a* KO mESCs (Fig. 6a). We reasoned that these newly formed insulation sites could interfere with repressive ultra-long loops between shared PRC1/PRC2 target genes. To explore this scenario, we analyzed the distances between newly formed insulation sites and Polycomb target genes. Strikingly, new insulation sites are located significantly closer to upregulated shared PRC1/PRC2 target genes than to all other classes of Polycomb and non-Polycomb genes (Fig. 6b). These data suggest that upregulation of a subset of shared PRC1/PRC2 target genes results from proximal changes in cohesin-mediated 3D chromosome architecture that cause a loss of ultra-long Polycomb loops. One prominent example of such cohesin-dependent dysregulation is the insulation gain region located between *Dlx2* and the *HoxD* gene cluster (Fig. 6c). In wildtype mESCs, *Dlx2* forms strong interactions traversing approximately 3 Mb with the *HoxD* gene cluster. Upon PDS5A deletion, these long-range interactions are lost and *Dlx2* expression is upregulated by more than 8-fold (Fig. 6c and Fig. 2e), yet PcG protein occupancy and associated histone modifications are either unaffected or only marginally reduced at *Dlx2* and the *HoxD* gene cluster (Fig. 5c and Fig. 3c, d). Instead, PDS5A deletion leads to a gain in insulation with increased cohesin binding near the *Sp9* gene, which is located between *Dlx2* and the *HoxD* gene cluster. Virtual 4-C viewpoints from the insulation gaining region (v2), as well as from *Dlx2* (v1) and the *HoxD* gene cluster (v3) reveal increased longer-range contacts in *Pds5a* KO mESCs, consistent with aberrant extension of chromosomal loops (Fig. 6c). This shift towards longer-range interactions is captured in Hi-C matrices as strengthening of two domains in between *Dlx2* and the *HoxD* gene cluster (dashed triangles) (Fig. 6c). We speculate that aberrant extension of chromosomal loops and strengthening of ectopic interaction domains comes at the expense of the ultra-long Polycomb loop between *Dlx2* and the *HoxD* gene cluster. A similar example showcasing how insulation gain might interfere with Polycomb long-range interaction and gene repression is represented by contacts between Hoxb13 and Ppp1r1b (Extended Data Fig. 6b).

Taken together our results argue that Polycomb loops are critical for silencing a subset of shared PRC1/PRC2 target genes. *Pds5a* KO causes changes in cohesin-dependent 3D genome architecture that perturb competing loops mediated by cPRC1. Notably, our data demonstrate that the resulting loss of Polycomb loops is linked to loss of silencing despite the presence of large repressive Polycomb chromatin domains. Thus, Polycomb repression takes place in a delicate spatial equilibrium with cohesin-dependent nuclear architecture, that is essential to maintain robust silencing at shared PRC1/PRC2 target genes.

## DISCUSSION

Here, we used a CRISPR-mutagenesis screen to identify novel regulators of cPRC1-induced gene silencing which revealed the cohesin regulatory subunit PDS5A. Subsequent KO in mESCs confirmed that PDS5A is required for repression of a subset of canonical Polycomb target genes. Notably, loss of developmental gene silencing is mostly uncoupled from changes in Polycomb repressive chromatin domains. Instead, PDS5A loss affects cohesin-dependent genome architecture which in turn perturbs competing loops mediated by cPRC1. Hence, our results strongly argue that ultra-long Polycomb loops are critical for robust silencing at a subset of canonical Polycomb target genes.

Intriguingly, even though the screen was performed in the absence of Dox, some of the genes that were identified (*Ring1b, Suz12, Pds5a*) are required for reporter gene silencing in the Polycomb *in vivo* assay only following Dox-dependent release of TetR-CBX7. We infer that the screen, which involved repeated FACS, relied on genes involved in the initiation or maintenance of reporter silencing. Overall, we predict that the epigenetic inheritance of reporter silencing involves a repressive Polycomb chromatin domain that engages in ultra-long interactions with shared PRC1/2 target genes. By causing cohesin dysfunction, PDS5A loss would disrupt the spatial integration of the silenced reporter locus from the existing network of Polycomb loops.

Previous reports have linked gene mutations of cohesin subunits to defects in Polycomb-dependent gene silencing, but the mechanisms had remained unclear. For example, a genetic screen for dominant suppressors of Polycomb-dependent silencing in *Drosophila* revealed several mutants in the *wapl* gene ^61^. In mammals, WAPL is a binding partner of PDS5A/B and the major regulator of cohesin off-loading and positioning ^37, 42, 56^. *Pds5a* KO mice exhibit developmental abnormalities, including skeletal malformations ^62^, that resemble the patterning defects of Polycomb mutant mice ^24, 63, 64^.

The capacity of cPRC1 to form 3D chromatin interactions that contribute to gene silencing has been previously demonstrated in *Drosophila* ^21, 28^. However, contributions to gene silencing by cPRC1 in mammalian systems has been controversial ^18, 19, 29, 30, 65, 66^. The repertoire of divergent PRC1 and PRC2 complexes increases dramatically from fly to mammals. Emerging hierarchies of Polycomb signaling cascades put vPRC1 and H2AK119ub front and center of Polycomb repression (reviewed in ^17^). Meanwhile, the repressive capacities of canonical pathways via PRC2, H3K27me3 and cPRC1 have been questioned, as Pcgf2/4-containing cPRC1 complexes have been shown to be largely dispensable in mESCs ^18, 19, 67^. Nevertheless, the repressive capabilities of cPRC1 have been clearly demonstrated ^29, 31, 66, 68, 69^.

Multiple studies in mammals have confirmed that cPRC1 mediates formation of chromatin loops underlying a 3D nuclear network of Polycomb target genes ^12, 22, 27, 70, 71^. Although the exact mechanisms that mediate these interactions remain unknown, PHC proteins, exclusively found in cPRC1, have been proposed to mediate these distal interactions via oligomerization of their SAM-domains ^24, 25^. Importantly, recent studies showed that cPRC1-mediated distal interactions and compaction of Polycomb domains are interdependent with genome organization by cohesin ^26, 44, 4426^. STAG2-containing cohesin complexes have been shown to contribute to compaction of Polycomb chromatin domains ^44^ and in Ewing sarcoma cells STAG2 depletion leads to a reduction in H3K27me3 and deregulation of PRC2 target genes ^72^. Furthermore, acute depletion of cohesin in mESCs resulted in stabilization of Polycomb chromatin loops and enhancement of transcriptional repression. These reports support a model that cohesin-mediated chromosome structure in the interphase nucleus normally restricts the formation of Polycomb looping ^26^. Here, we discovered that ultra-long Polycomb loops are preferentially affected by cohesin dysfunction caused by *Pds5a* KO. This finding is consistent with overarching principles in nuclear organization: compartmental domains form between similar chromatin states, leading to the checkerboard pattern observed in Hi-C data, where A and B compartments tend to interact with other A and B compartments rather than A with B or vice versa. Cohesin- and CTCF-dependent loop formation on the one hand facilitates interaction frequencies within identical compartmental domains, but on the other hand loops across TADs of A and/or B compartments restrict their segregation. Thus, CTCF depletion removes this restriction of segregation, leading to increased compartmentalization despite a loss of loops ^42, 73–76^. In addition, stabilizing cohesin enhances loop extrusion and increases the restriction of compartment segregation ^37, 42, 75^. Hence, this model could explain why PDS5A loss preferentially affects longer loops, that might be formed by such natural interaction of similar chromatin domains over very long distances that usually are not disturbed by cohesin loop extrusion ^75^.

Nevertheless, to our knowledge a causal link between Polycomb-mediated long-range interactions and repression of Polycomb target genes has not been described. Polycomb target genes are generally located within the active (A) compartment in the nucleus of mESCs and thereby in spatial proximity to actively transcribed genomic regions ^77^. We speculate that Polycomb-dependent silencing of canonical target genes involves multiple parallel mechanisms including repressive histone modifications and long-range interactions. By uncoupling loss of Polycomb silencing and loop interactions from changes in repressive chromatin modifications, our data argue that 3D organization by itself has repressive function potentially by tethering PRC1/PRC2 target genes away from transcriptional co-activators and/or RNA Polymerase. Therefore, when combined with repressive chromatin modifications, which promote Polycomb feedback mechanisms, spatial aggregation by Polycomb loops would effectively enhance robust gene silencing.

The relative contribution of each of these mechanisms is likely locus-specific and may vary in different cell types. For example, alternative incorporation of paralogous cPRC1 subunits, such as CBX proteins or PHC proteins may influence the specific regulation of Polycomb 3D network formation as a mechanism of repression. Hence, future studies are needed to discern how cPRC1 and Polycomb-dependent genome architecture control target gene silencing in the context of different cell types and in disease.

## Acknowledgements

We are grateful to all members of the Bell, Farnham and Jadhav laboratories, as well as Alexander Stark, Martin Leeb, Jan Michael Peters, Gordana Wutz, Diana Hargreaves, Jesse Dixon, Suhn Rhie and Geoffrey Fudenberg for feedback and discussions. We especially thank Ilya Flyamer for sharing Hi-C protocols and advice on Hi-C data analysis. We thank Jan Michael Peters and Gordana Wutz for sharing antibodies. We thank the Vienna Biocenter Core Facility Next Generation Sequencing. The GMI/IMBA/IMP Scientific Service units and the BioOptics facility. We thank Life Science Editors for editorial assistance.

## Funding

O.B. and U.E. were supported by the Austrian Academy of Sciences. O.B. was supported by the New Frontiers Group of the Austrian Academy of Sciences (NFG-05), the Human Frontiers Science Programme Career Development Award (CDA00036/2014-C), and start-up funding from the Norris Comprehensive Cancer Center at Keck School of Medicine of USC.

### Author contributions

D.B., H.F.M., O.B. initiated and designed the study. D.B., H.F.M. generated cell lines. R.Y. generated parental cell lines. U.E., G.M. provided the CRISPR sgRNA library and helped with the screen design. D.B., H.F.M. performed CRISPR-Cas9 genetic screen. J.W. and G.M. analyzed CRISPR-Cas9 genetic screen data. D.B. performed molecular biology, RNA-seq, ChIP-seq and Hi-C experiments. S.G., Q.Z. participated in experiments. RNA-seq, ChIP-seq and Hi-C data analysis was conducted by D.B. O.B. supervised all aspects of the project. The manuscript was prepared by D.B. and O.B.. All authors discussed results and commented on the manuscript.

### Data, Material and Code availability

All NGS data reported in this study has been deposited at the Gene Expression Omnibus (GEO) database under accession number GSE *********

## METHODS

### Cell lines

All cell lines used directly in this study or for generating mutants were diploid mESCs derived from originally haploid HMSc2 termed AN3-12 ^45^. TetR-CBX7 reporter mESCs with 7x TetO DNA binding sites flanked by GFP and BFP reporter genes were previously described ^31^. Pds5a gene-trap (GT) KO and its corresponding wild type mESCs with genetrap insertion in the non-disruptive orientation were aquired from the Haplobank repository (Cell IDs: 10388IH and 10388MH) ^45^.

### Cell culture conditions

All mESCs were cultivated without feeders in high-glucose-DMEM (Corning 10-013-CV) supplemented with 13.5% fetal bovine serum (Corning 35-015-CV), 10 mM HEPES pH 7.4 (Corning, 25-060-CI), 2 mM GlutaMAX (Gibco, 35050-061), 1 mM Sodium Pyruvate (Corning 25-000-Cl), 1% Penicillin/Streptomycin (Sigma, P0781), 1X non-essential amino acids (Gibco, 11140-050), 50 mM β-mercaptoethanol (Gibco, 21985-023) and recombinant LIF. Cells were incubated at 37°C and 5% CO_2_ and were passaged every 48 hours by trypsinization in 0.25% 1x Trypsin-EDTA (Gibco, 25200-056). In order to reverse of TetR-CBX7 fusion protein binding 1 µg/ml Doxycycline (Sigma, D9891) was added to cell culture medium.

### Generation of CRISPR-Cas9 mutants

TetR-CBX7 RING1B and PDS5a loss of function (LOF) as well as An3-12 Pds5a LOF mutants were generated using CRISPR-Cas9 technology. sgRNAs targeting Pds5a (5’-TGTCTCTGCAGAGTGGAACG-3’) or Ring1b (5’-ACAAAGAGTGTCCTACCTGT -3’) were introduced into a modified version of the vector plasmid pX330-U6-Chimeric_BB-CBh-hSpCas9 (Addgene #42230) that yields a BFP marker for selection (Gift by J. Zuber). Plasmid transfection was achieved by electroporation with the NEON transfection system (Invitrogen, MPK5000). 36 hours post transfection cells were FACS sorted for Cas9-BFP and 1000-2000 cells seeded for clonal expansion on a 15cm plate. 7-10 days later colony forming clones were individually picked, the targeted loci genotyped and loss of function confirmed by western blot.

### Hoechst-staining

For cell cycle profiling mESCs were trypsinized and genomic DNA was stained with Hoechst 33342 (20 mM; Thermo Fisher Scientific Cat. # 62249) for 30 min at 37°C and 5% CO_2_. Hoechst immunofluorescence was measured by flow cytometry on a FACSAria II cell sorter (BD Biosciences).

### AP-staining

One thousand cells were seeded and grown to form colonies at low density on 15 cm tissue culture dishes for 7 days. On day 7, dishes were washed with 100 mM tris (pH 8) and AP activity assay was performed using the VECTOR Blue AP Substrate Kit (Vector Laboratories, VECSK-5300) according to the manufacturer’s instructions. Following AP staining, stained colonies were fixed in 4% formaldehyde overnight. Plates were rinsed with 1x PBS the following day and images taken on a brightfield microscope (EVOS XL Core system).

### Pluripotency marker staining

To assess pluripotency of PDS5a loss of function mutants, we applied intracellular staining of OCT3/4, SOX2 and SSEA1. Single cell suspensions of wild type and Pds5a KO mESCs were permeabilized and fixed using the fixation/permabilization buffer (R&D systems), washed twice with 1x PBS and stained using the H/MM pluripotent Stem Cell Multi-Color Flow Cytometry kit (R&D Systems) according to vendor’s protocol. Flow cytometry data was collected on an Attune NxT equipped with Attune NxT v3.1 acquisition software. Final data analysis was performed using FlowJo (10.7.1).

### Western blot

10 million mESCs were subsequently lysed in Buffer A (25 mM Hepes pH 7.6, 5 mM MgCl_2_, 25 mM KCl, 0.05 mM EDTA, 10% Glycerol, 1 mM DTT, 1 mM PMSF, 1× Complete Mini protease inhibitor, Roche) resuspended in RIPA buffer (150 mM NaCl, 1% triton, 0.5% sodium deoxy-cholate, 0.1% SDS, 50 mM Tris pH 8.0). Lysates were homogenized by sonication using a Bioruptor Pico (Diagenode) and concentration determined by Bradford assay (Biorad). 4x non-reducing Laemmli SDS sample buffer (Alfas Aesar, #J63615AD), 10 mM final DTT and 0.5% final BME were added to 20 µg total protein/sample and boiled at 95°C for 5 min. Samples were separated on NuPAGE 4– 12% Bis-Tris gels (Invitrogen) in Bis-Tris running buffer (Novues Biologicals) and transferred on a Merck Chemicals Immobilon-FL Membrane (PVDF 0.45 µm). After blocking the membranes (5% non-fat dry milk in 1× PBS, 0.1% Tween 20) the blots were incubated o/n with the primary antibodies in 5% non-fat dry milk in 1× PBS and 0.1% Tween 20. Antibodies used: PDS5a (Millipore Sigma #SAB2101764) 1:1000; RING1B (Cell Signaling D22F2) 1:1000; SMC3 (Bethyl Laboratories A300-060A) 1:2000; SUZ12 (Cell Signaling D39F6) 1:1000; LAMIN B1 (Abcam ab16048) 1:15000; H2AK119ub (Cell Signaling D27C4) 1:20000; H3K27me3 (Diagenode p069-050) 1:1000. Next, the membranes was incubated with corresponding secondary IRDye 800CW Goat anti-Rabbit IgG (H+L) (LICOR) or IRDye 680RD Goat anti-Mouse IgG (H+L) (LICOR) antibodies and imaged on an Odyssey CLx Near-Infrared Imaging System (LICOR).

### Genetic CRISPR-Cas9 screen

For the genetic CRISPR-Cas9 mutagenesis screen, EF1a promoter driven hSpCas9 with a hygromycin resistance marker (modified version of Addgene #52961) was stably integrated via lentiviral transduction into the previously described TetR-CBX7 reporter mESC cell line, which contains 7x TetO DNA binding sites flanked by GFP and BFP reporter genes ^31^.

For CRISPR-Cas9 mutagenesis, a sgRNA library targeting 6,560 nuclear factors with 4 sgRNAs per gene and 112 nontargeting controls was utilized (described in ^32^. For retroviral library generation, the barcoded plasmid library of sgRNAs, containing neomycin resistance for selection, was packaged in PlatinumE cells (Cell Biolabs) according to the manufacturer’s recommendations.

300 million TetR-CBX7 reporter mESCs were infected with a 1:10 dilution of the harvested virus-containing supernatant PlatinumE cell medium for 24 hrs in the presence of 2 μg/ml polybrene (Santa Cruz Biotechnology, SACSC-134220). The 300 million cells were divided into 3x 100 million sets (10x 15 cm plates of 10 million cells each) that were treated as 3 separate replicates throughout entirety of the mutagenesis screen and sgRNA NGS sequencing. 24 hrs post infection, neomycin-resistance selection was started on the infected cells by addition of G418 (Gibco) at 0.5 mg/ml. After 24 hrs of selection, each replicate was expanded from 10 15-cm dishes to 20 dishes.

Subsequently, for the duration of the neomycin selection cells were always maintained at a minimum of 300 million cells. After 5 days and completion of G418 selection, half of the cells were cultured in mESC medium containing doxycycline for 3 days and the other half without doxycycline. GFP positive cell populations of both doxycycline treated and untreated populations were sorted on a FACSAria II cell sorter (BD Biosciences) and flow cytometry data analyzed with FlowJo software. Unsorted mutant populations were served as background controls. Genomic DNA was isolated from GFP-positive sorted and unsorted cells, their sgRNA cassettes amplified by PCR and subjected to NGS sequencing on an Illumina HiSeq 2500. Data analysis was performed as previously described in (Michlits 2017) and gene enrichment determined using MAGeCK ^33^.

### RNA-seq

5 million mESCs were trypsinized and collected by centrifugation. Resulting cell pellets were washed in 1x PBS and resuspended in 1x DNA/RNA protection reagent (Monarch Total RNA Miniprep Kit, NEB). Subsequently, cells were lysed and total RNA extracted following the mammalian cell protocol including optional on-column DNase I treatment. For RNA-seq library preparation, 1 µg of total RNA per sample was enriched for poly-A using the NEBNext Poly(A) mRNA Magnetic Isolation Module (NEB, E7490) and final RNA-seq libraries generated using the NEBNext Ultra II Directional RNA Library Prep kit (NEB, E7760 and NEBNext Multiplex Oligos (NEB, E7335/E7500). Final libraries were sequenced as 150 bp paired-end reads on the Illumina HiSeq platform.

### RNA-seq Data Analysis

Raw paired-end RNA-seq reads were aligned to the mm10 genome using STAR-2.6.1c^78^. Overlap of STAR-aligned reads with genes was performed using HTSeq count function ^79^ with stranded=reverse option and the GRCm38 version 94 GTF file. The HTseq count matrix was pre-filtered to exclude genes with a read count below 10. Differential gene expression analysis was performed using DESeq2 ^80^ using the “apeglm” method ^81^ for LFC shrinkage. We applied a threshold of p-adj < 0.05 and fold change > 0.5 or -0.5 for gene expression changes to be considered significant. Visualization of RNA-seq data was performed using custom R scripts and ggplot2. Gene ontology analysis for significantly deregulated genes was performed using custom R scripts and clusterProfiler reference ^82, 83^.

### Calibrated ChIP-seq (cChIP) and ChIPCap-seq

30 million mESCs and HEK293T cells were collected, washed once in 1x PBS and crosslinked for 7 min in 1% formaldehyde. The crosslinking was quenched by addition of 125 mM glycine and incubated on ice. The crosslinked cells were pelleted by centrifugation for 5 min at 1200g at 4 °C. Nuclei were prepared by washes with NP-Rinse buffer 1 (10 mM Tris pH 8.0, 10 mM EDTA pH 8.0, 0.5 mM EGTA, 0.25% Triton X-100) followed by NP-Rinse buffer 2 (10 mM Tris pH 8.0, 1 mM EDTA, 0.5 mM EGTA, 200 mM NaCl). Afterwards, the nuclei were washed twice with shearing buffer (1 mM EDTA pH 8.0, 10 mM Tris-HCl pH 8.0, 0.1% SDS) and subsequently resuspended in 900 µL shearing buffer with added 1× protease inhibitors complete mini (Roche). Chromatin was sheared by sonication in 15 ml Bioruptor tubes (Diagenode, C01020031) with 437.5 mg sonication beads (Diagenode, C03070001) for 6 cycles (1 min on/1 min off) on a Bioruptor Pico sonicator (Diagenode). For each ChIP reaction 4 % HEK293T-derived human spike-in lysate was combined with mESC lysate and incubated in 1x IP buffer (50 mM HEPES/KOH pH 7.5, 300 mM NaCl, 1 mM EDTA, 1% Triton X-100, 0.1% DOC, 0.1% SDS), with following appropriate antibodies at 4 °C o/n a rotating wheel: H3K27me3 (Diagenode, C15410195), RING1B (Cell Signaling, D22F2), PDS5a (Millipore Sigma #SAB2101764), SUZ12 (Cell Signaling D39F6), H2AK119ub (Cell Signaling D27C4), PCGF1 (Abcam ab202395), RAD21 (Abcam ab992), CTCF (Millipore 070729).

Antibody-bound chromatin was captured using Dynabeads protein G beads (Thermofisher #10004D) for 4 hours at 4°C. ChIP washes were performed as described previously ^51^. ChIPs were washed 5x with 1x IP buffer (50 mM HEPES/KOH pH 7.5, 300 mM NaCl, I mM EDTA, 1% Triton-X100, 0.1% DOC, 0.1% SDS), or 1.5x IP buffer for H3K27me3 and H2AK119ub, followed by 3x washes with DOC buffer (10 mM Tris pH 8, 0.25 mM LiCl, 1 mM EDTA, 0.5% NP40, 0.5% DOC) and 1x with TE/50 mM NaCl. ChIP DNA was eluted 2x in elution buffer (1% SDS, 0.1 M NaHCO3) at 65°C for 20 min, RNase A treated for 30 min at 37 °C, Proteinase K treated for 3 hrs at 55 °C and crosslinks were reversed o/n at 65 °C. The following day, ChIP samples and corresponding inputs were purified by PCI extraction and DNA precipitation.

### cChIP-seq and ChIPCap-seq library preparation

Libraries were prepared using the NEXTflex ChIP-Seq kit (Bio Scientific) following the “No size-selection cleanup” protocol. Libraries were purified using Agencourt AMPure XP (Beckman Coulter) and amplified using the KAPA Real-Time Library Amplification Kit (KAPABiosystems) following the manufacturer’s instructions.

ChIPCap-seq libraries were prepared identically to ChIP-seq libraries. After PCR amplification the libraries were enriched for loci of interest using the MYbaits kit DNA capture target enrichment system (Arbor Biosciences) according to manufacturer’s manual. 120 nucleotides long MYbaits sequence capture probes (Arbor Biosciences) were custom designed against 25 mm9 genomic loci (Supplementary Table 3).

Library quality control including determination of average size and concentration was performed prior to sequencing by commercial Next Generation Sequencing providers. NGS libaries were eventually sequenced as 150 bp paired-end reads on the Illumina HiSeq platform.

### ChIP-seq Data Analysis

Raw reads were mapped to the custom concatenated mouse (mm10) and spike-in human (hg38) genome sequences using bowtie 2 with “–no-mixed” and “no-discordant” options (Langmead and Salzberg, 2012). Subsequently, low quality reads were filtered using SAMtools ^84^, duplicated reads were discarded with the Picard toolkit (http://broadinstitute.github.io/picard/) and only unique mapped reads were retained.

For visualization uniquely mapped mouse reads were normalized by random subsampling with samtools using calibration factors calculated from the corresponding hg38 spike-in reads as described previously ^19, 85, 86^. High correlation between replicates was confirmed using multiBamSummary and plotCorrelation functions from deepTools^87^ before merging for visualization and downstream analysis. Genome coverage tracks (bigWig files) were produced with MACS2’ pileup function ^88^ and heatmaps and profile plots generated with deepTools ^87^.

Peaks were called on each replicate independently using MACS2 ^88^. Peaks overlapping with a custom-build blacklist were discarded to remove sequencing artifacts and only peaks called in both replicates were retained for downstream analysis.

### In Situ Hi-C

Hi-C was performed as previously described ^77^ with modifications described in ^22^ as following: 5 million mESCs were crosslinked in 1% formaldehyde for 10 mins before the reaction was quenched by adding 0.2 M final glycine. Cells were permeabilized in lysis buffer (0.2% IGEPAL, 10 mM Tris-HCl pH 8.0, 10 mM NaCl, 1x Halt Protease inhibitor cocktail) and nuclei isolated in NEBuffer 3 supplemented by 0.3% SDS at 62 °C for 10 min. SDS was quenched with 1% Triton X-100 at 37°C for 60 min, the nuclei pelleted and resuspended in 250 µl of 1x DpnII buffer with 600 U DpnII (NEB). Following o/n digestion at 37°C, 200 U were added for 2 hrs. DpnII was inactivated for 20 min at 65°C before the DNA ends were filled-in and biotin-marked using Klenow, d(C/G/T)TPs and biotin-14-dATP for 90 min at 37°C. Proximity ligation was performed using T4 DNA ligase (NEB) for 4 hrs at room temperature. Subsequently, nuclei were spun down, resuspended in 200 µl mQ water and digested with proteinase K for 30 min at 55°C in presence of 1% SDS. For crosslink reversal 1.85 M final NaCl was added and samples incubated at 65°C o/n. The next day, sample DNA was ethanol precipitated and sheared in 500 µL sonication buffer (50 mM Tris pH 8.0, 0.1% SDS, 10 mM EDTA) on a Bioruptor Pico sonicator (Diagenode). DNA was then concentrated on Amicon ultra 0.5 30K filter units (Millipore), biotin-pulldown performed using MyOne Streptavidin T1 beads (Life technologies, 65602) and used for NGS library preparation. DNA ends were repaired, biotin removed from unligated ends and NEXTFLEX DNA barcoded adapters (Perkin Elmer) were ligated. Desired PCR cycle numbers were determined in test endpoint PCRs using Q5 DNA polymerase (NEB M0491L). Final HiC libraries were generated from 4-6 individual PCR reactions, which were pooled and subjected to cleanup and size-selection using AMPure beads (Beckman Coulter A63882). Samples were first test sequenced to check library quality before selected Hi-C ibraries were sequenced at greater depth (Supplementary Table 2).

### Hi-C data analysis

Hi-C data were analyzed using the HiC-Pro (2.11.1) pipeline ^89^. Read mapping to the mm10 genome was performed using bowtie 2 ^90^ within the HiC-Pro pipeline. PCR and optical duplicates as well as reads with MAPQ < 30 were removed. Filtered valid HiC contact data was binned and raw and ICE normalized .hic contact matrices were generated. We also produced balanced single and multi-resolution .cool and .mcool cooler files for visualization in HiGlass. Virtual 4C tracks were obtained using the hicPlotViewpoint function of HiCExplorer ^91–93^. For reference the same analysis was applied to published deep Hi-C data from mESCs ^59^. To obtain wild type mESC TAD and loop information we applied juicer tools ^94^ “arrowhead” and “HiCCUPS” to the Bonev 2017 mESC dataset. To delineate A and B compartment information, “Eigenvector” function of juicer tools was applied to 250kb wildtype and Pds5a KO mESC data created in this study.

For pileup analysis at HiC loops we used coolpup.py ^95^ and took averaged the calculated observed over expected interactions within a 105 kb x 105 kb window centered on the loops at 5 kb resolution.

For calculation and visualization of compartment strengths, relative contact probabilities, insulation scores and differential HiC contact matrices we used the R package GENOVA (https://github.com/robinweide/GENOVA) ^96^. Insulation score analysis was conducted as previously described in (^97^.Insulation scores ^60^ were computed at 10kb resolution using GENOVA. Bins with an insulation score greater than -1 were excluded from the analysis. To qualify as differentially insulated bins, either wild type or Pds5a KO mESCs had to be lower than - 0.2 to exclude very lowly insulated portions of the genome. Insulation gaining and losing regions were defined as bins that had an absolute change in insulation between wild type and Pds5a KO of 0.2.

## EXTENDED DATA FIGURE TITLES AND LEGENDS

**Extended Data Fig. 1.**
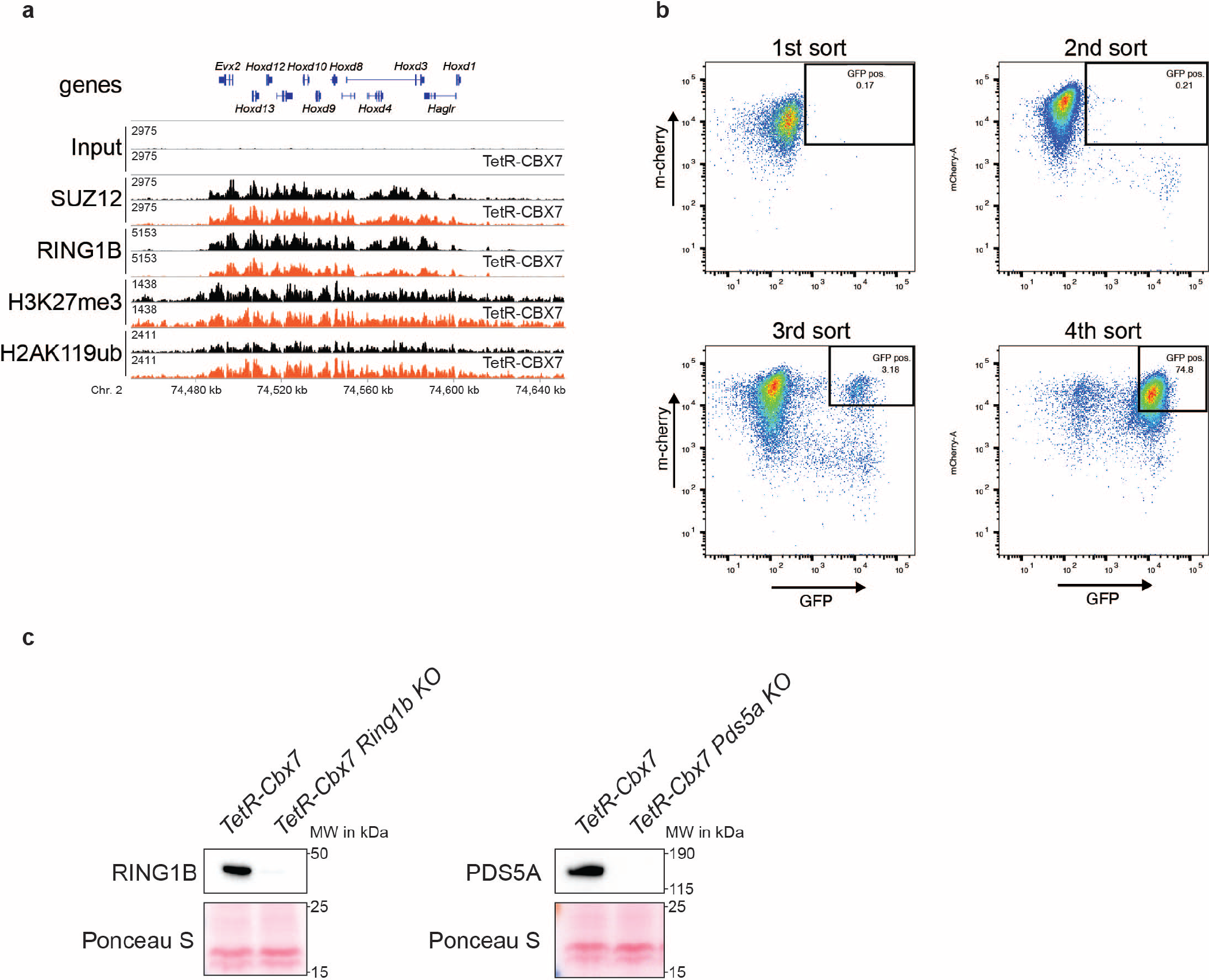
CRISPR screen of cPRC1-dependent gene silencing reveals PDS5A. **a,** Genomic ChIP-CapSeq screenshot of polycomb proteins and histone modifications before (black) and after TetR-Cbx7 expression (orange) at HoxD cluster. **b,** Flow cytometry scatterplots of CRISPR screen sorting scheme to enrich for GFP-positive reporter mESCs. GFP expression of reporter is shown on x-axis and TetR-Cbx7 expression represented by m-cherry on y-axis. Sorting gates applied are indicated in each plot. **c,** Western blots for RING1B and PDS5A of wild type TetR-Cbx7 mESC reporter cell lines and *Ring1b* KO (left panel) and *Pds5a KO* (right panel) clonal cell lines.

**Extended Data Fig. 2.**
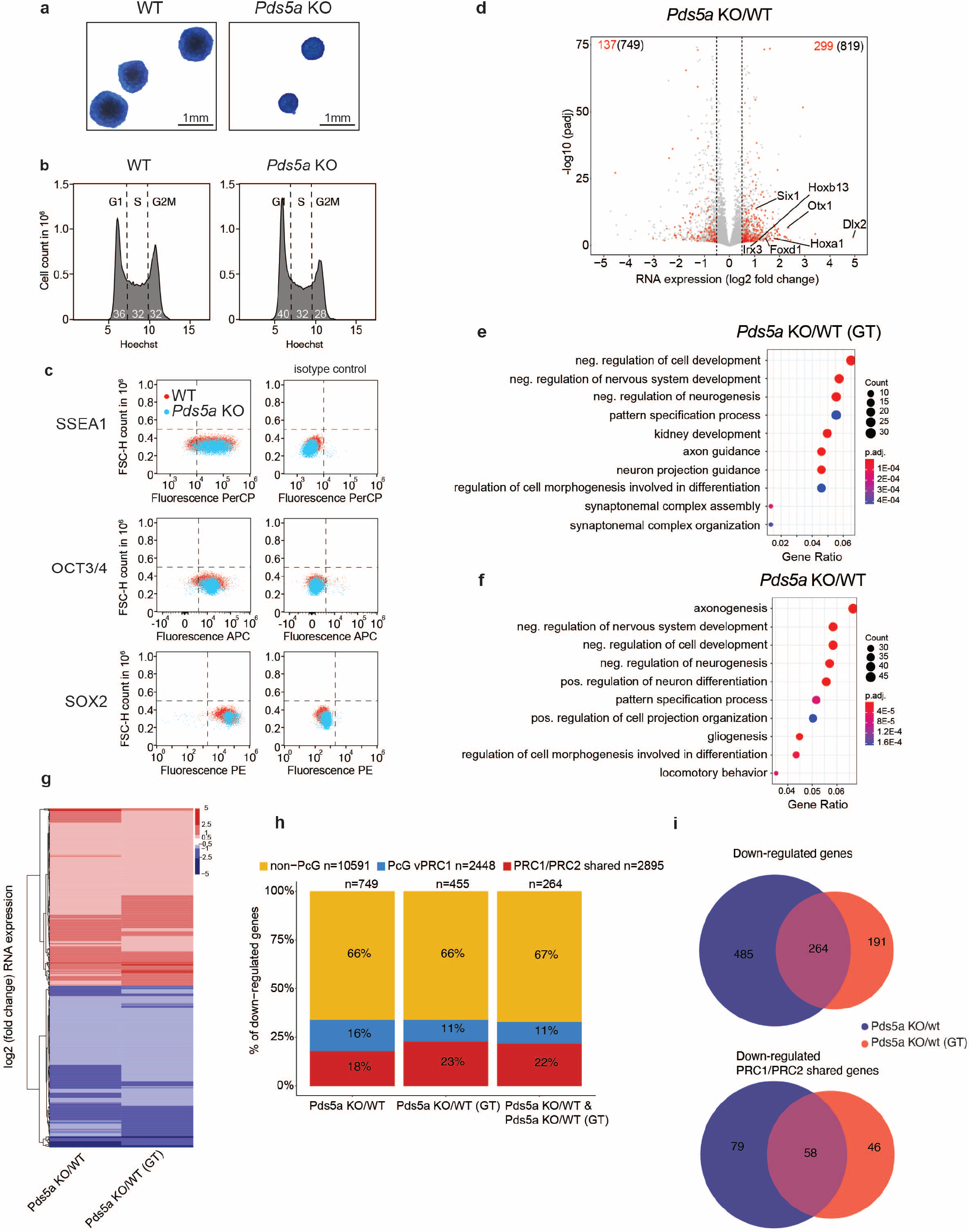
Pds5a KO does not impair mESC stem cell markers or cell cycle profiles and causes de-repression of canonical PRC1/PRC2 target genes in second independent Pds5a KO cell line. **a,** Alkaline phosphatase staining of wildtype and Pds5a KO mESCs. **b,** Flow cytometry histogram after Hoechst staining showing cell cycle distribution plot of G1, S and G2M phase in wild type and Pds5a KO. **c,** Flow cytometry scatterplots of stem cell marker staining (SSEA1, OCT3/4 and SOX2) including corresponding antibody isotype controls (right panels). **d,** Volcano plot of RNA-seq expression change in PDS5A CRISPR-Cas9 KO vs. wild type mESCs (Pds5a KO/WT). Genes from the PRC1/PRC2 shared class (n=2895) with increased or decreased expression are marked in red (padj < 0.05 and log2 fold change > 0.5 or < -0.5). Total numbers of up-/and down-regulated genes are indicated in the top left and right corners (PRC1/PRC2 shared class in red and of all genes in black). **e,** Gene ontology analysis plots of enriched gene groups among all up-regulated genes (padj < 0.05 and log2 fold change > 0.5) in PDS5A genetrap KO mESCs and **f,** PDS5A CRISPR-Cas9 KO mESCs. **g,** Heatmap of log2 fold changes in RNA-seq of all de-regulated genes (padj < 0.05 and log2 fold change > 0.5 or < -0.5) upon PDS5A loss for PDS5A genetrap (GT) and PDS5A CRISPR-Cas9 KO mESCs. **h,** Stacked bargraphs of percent gene distribution of down-regulated genes in both Pds5a KO cell lines individually and in their overlap. **i,** Euler diagrams of overlap of all (top), as well as only PRC1/PRC2 shared down-regulated genes (bottom) between both Pds5a KO cell lines.

**Extended Data Fig. 3.**
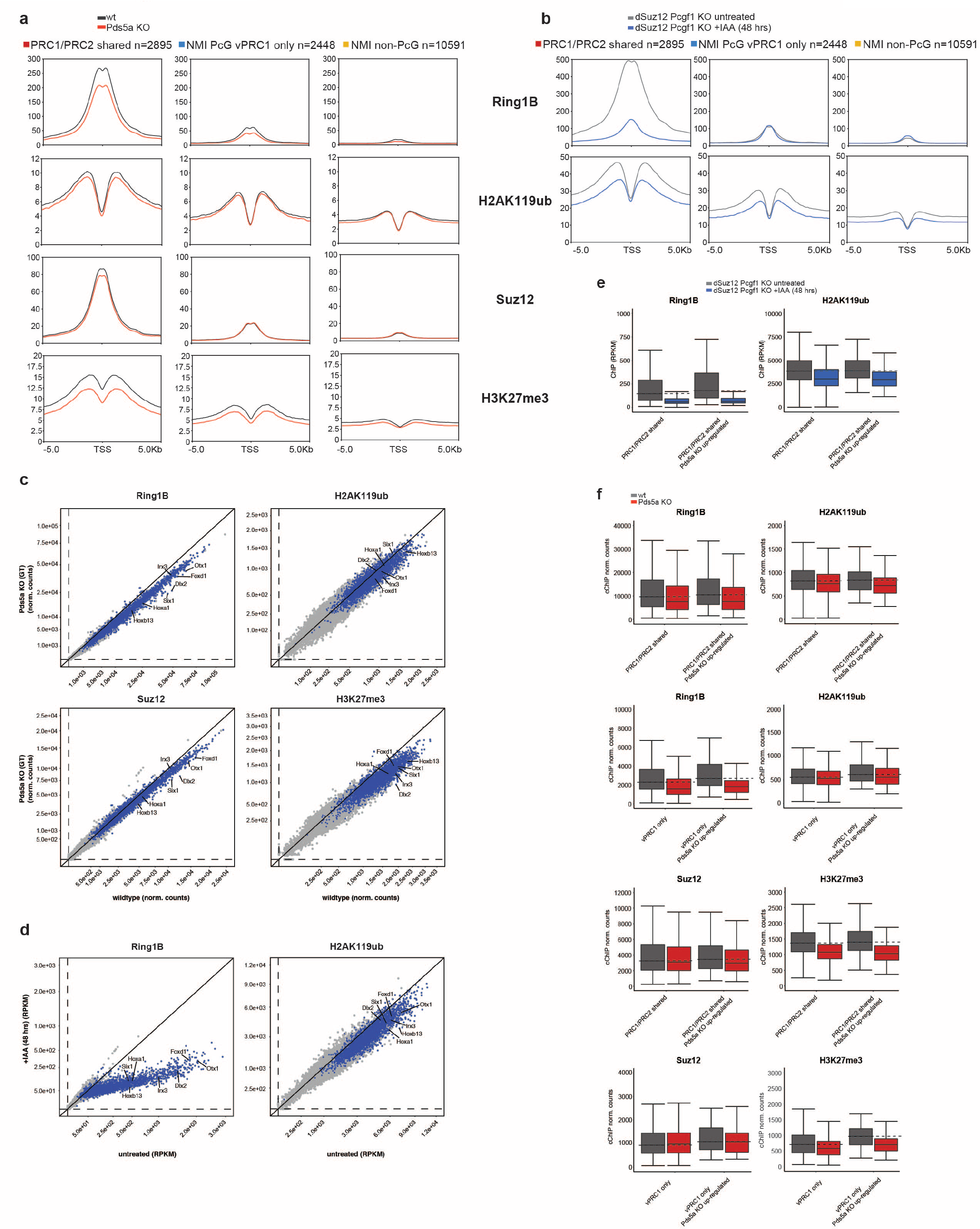
PDS5A deletion has minimal effect on Polycomb repressive chromatin domains. **a,** Profile plots quantifying mean cChIP signal for each IP (Ring1b, H2AK119ub, Suz12, H3K27me3) and gene class in Fig 3A for wildtype and Pds5a KO mESCs and **b,** 48 hrs IAA-treated vs. untreated dSuz12 Pcgf1 KO mESCs. **c,** Scatterplots of cChIP signal around TSS for each gene (Sum of TSS +/- 2.5 kb). Genes from the PRC1/PRC2 shared gene class are colored in red and selected representative genes are labeled individually (see Fig 2d) for wildtype and Pds5a KO mESCs and **d,** 48 hrs IAA-treated vs. untreated dSuz12 Pcgf1 KO mESCs. **e,** Boxplots of PRC1 and PRC2 complex and histone modification cChIP signal within the entire PRC1/PRC2 shared or vPRC1 only gene class and their corresponding subsets of transcriptionally up-regulated genes for 48 hrs IAA-treated vs. untreated dSuz12 Pcgf1 KO mESCs and **f,** wildtype and Pds5a KO mESCs.

**Extended Data Fig. 4.**
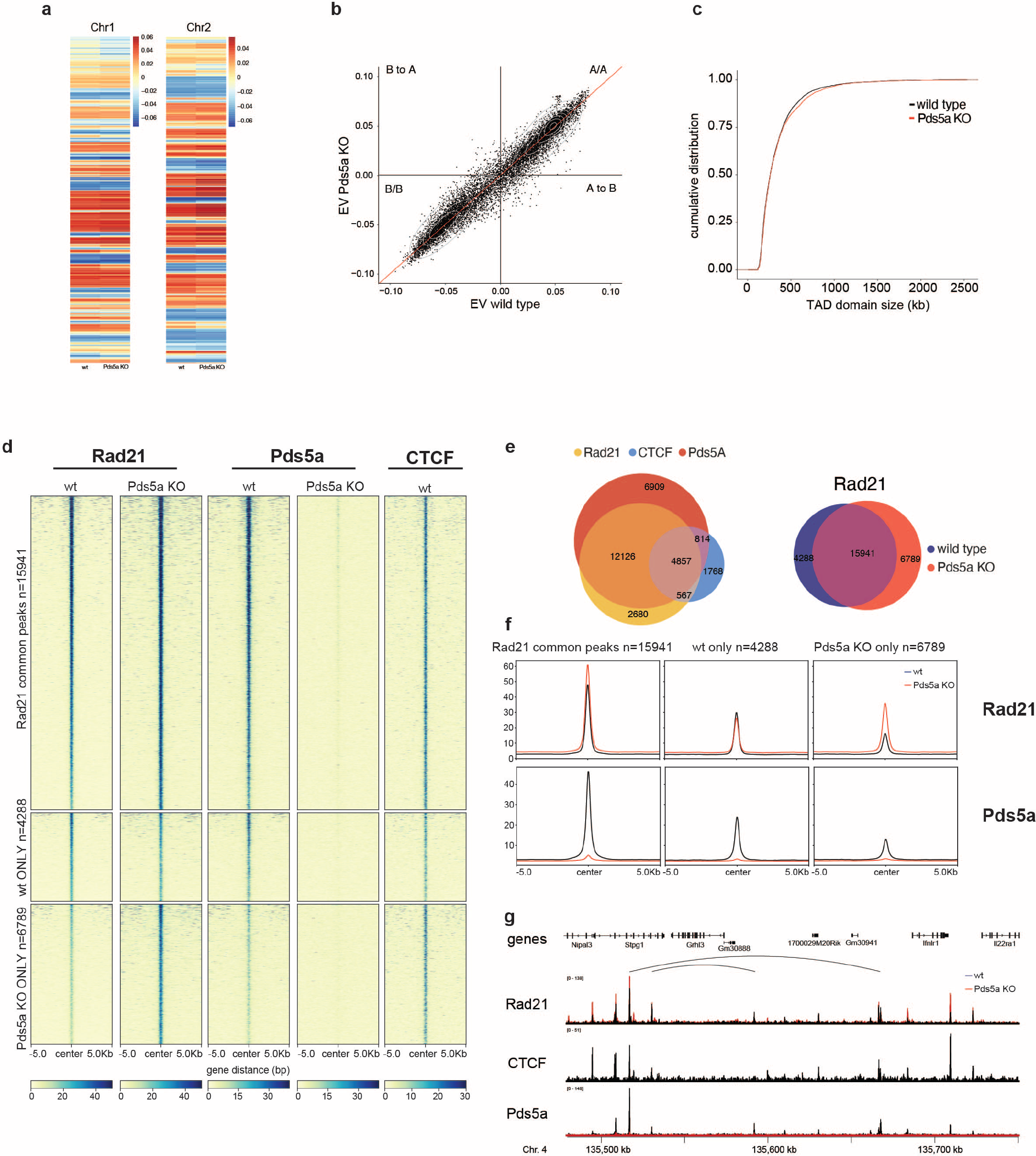
PDS5A colocalizes with cohesin and stabilizes chromatin binding. **a,** Heatmaps of compartment signal at 250kb resolution for chromosomes 1 and 2 in wildtype and Pds5a KO. **b,** Scatterplot of compartment signal (Eigenvector) for each 250kb bin in the genome in wildtype vs. Pds5a KO. **c,** Cumulative distributions of called TAD domains in wildtype and Pds5a KO. **d,** Wildtype and Pds5a KO cChIP-seq heatmaps of Rad21, PDS5A and CTCF. cChIP-seq signal is plotted around common wild type/KO (n=15941), wt only (n=4288) or Pds5a KO only (n=6789) peak centers +- 5kb. **e,** Euler diagrams of overlap of wild type CTCF, Rad21 and PDS5A peaks (left) and overlap of wild type and Pds5a KO called Rad21 peaks. **f,** Profile plots quantifying mean cChIP signal of Rad21 and PDS5A cChIP at Rad21 peaks. **g,** IGV genomic cChIP-seq screenshot illustrating CTCF (wild type only), Rad21 and PDS5A in wildtype and Pds5a KO.

**Extended Data Fig. 5.**
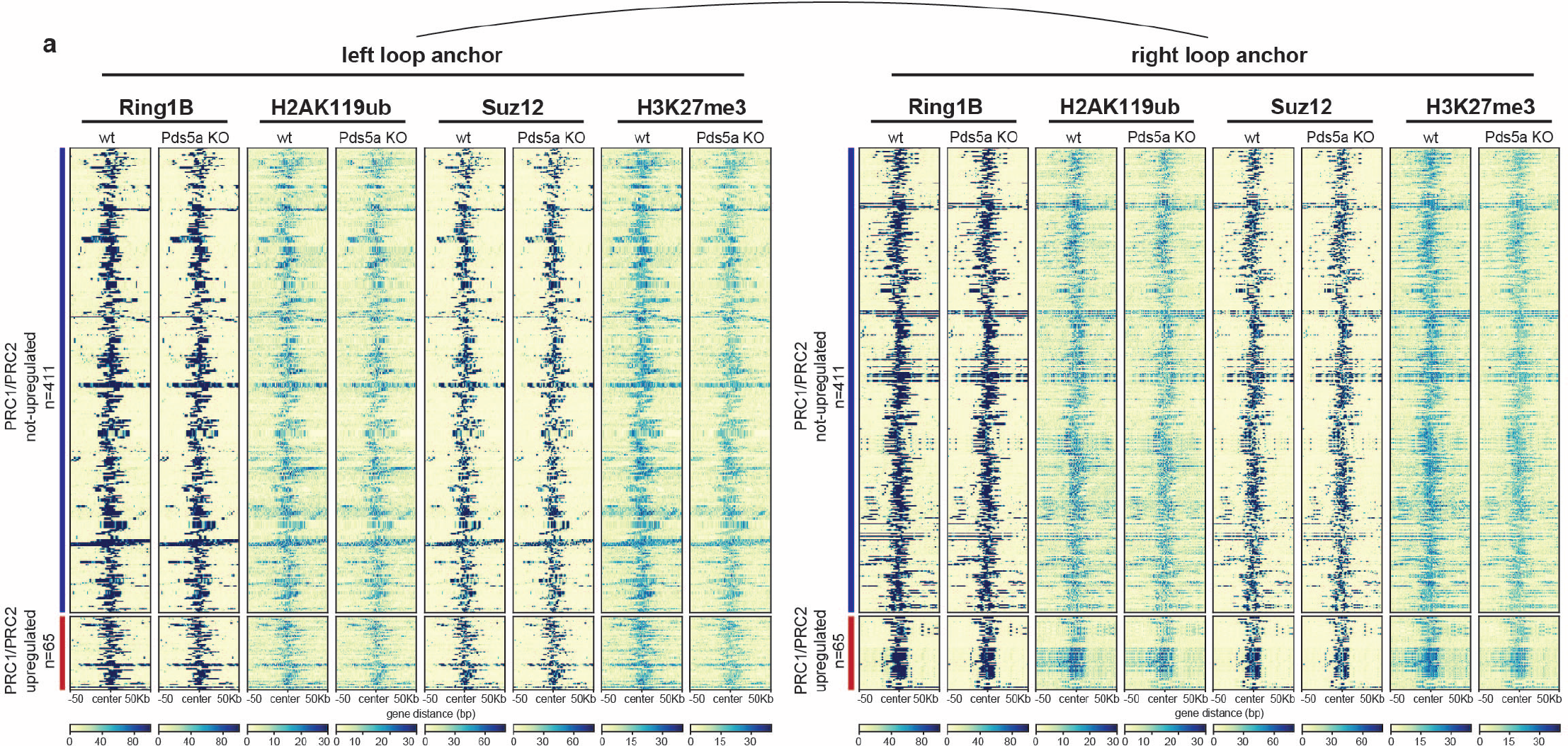
PDS5A is required for maintenance of repressive polycomb loops crossing ultra-long distances. **a**, cChIP-seq heatmaps of PRC1 (Ring1b and H2AK119ub) and PRC2 (Suz12 and H3K27me3) at Polycomb loops between shared PRC1/PRC2 genes. Heatmap is divided in loops between not-upregulated PRC1/PRC2 shared target genes (left and right loop anchors; n=411) and loops between upregulated PRC1/PRC2 shared target genes (left loop anchor represents the not upregulated side and right loop anchor the Pds5a KO transcriptionally upregulated side; n=65).

**Extended Data Fig. 6.**
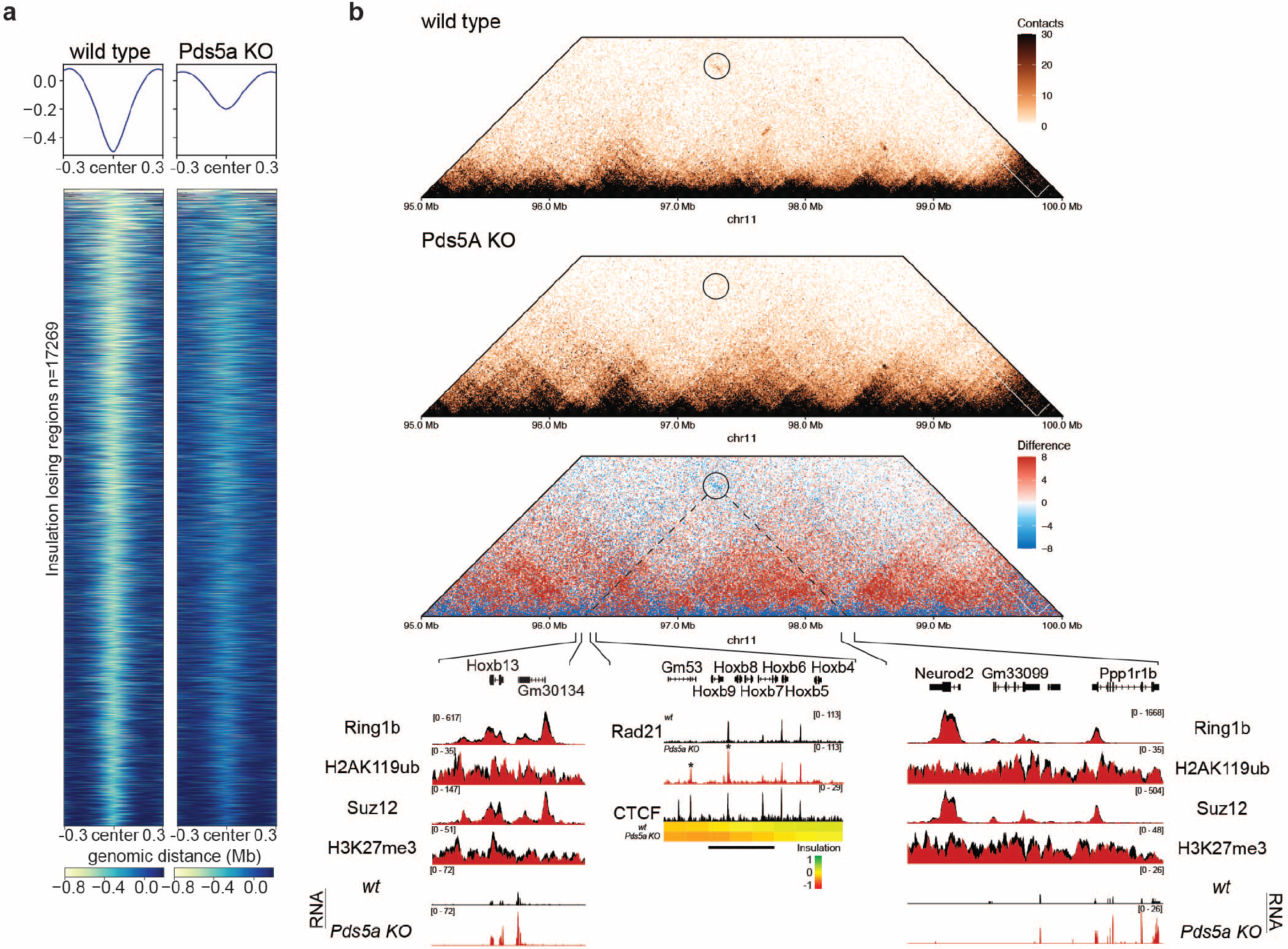
Loss of repressive Polycomb loops is linked to cohesin-mediated insulation gain. **a,** Heatmaps and average profile plots of insulation score of genomic 10kb bins (+/- 300kb) in wild type and Pds5a KO. Shown are insulation losing bins (n=17269; minimum insulation score increase of 0.2). **b,** Wild type and Pds5a KO HiC matrices and a differential HiC matrix (blue = loss; red = gain of contacts in Pds5a KO; 10kb resolution and ICE-normalized). Genomic screenshots of polycomb group proteins and histone modification cChIP are displayed for the HoxB cluster and Ppp1r1b. Insulation scores, as well as cChIP IGV screenshots of cohesin (Rad21) and CTCF are shown at the Insulation gaining region (indicated by black bar). Change in RNA-seq signal for Dlx2 is plotted underneath.

## REFERENCES

1. Kassis, J. A., Kennison, J. A. & Tamkun, J. W. Polycomb and Trithorax Group Genes in Drosophila. Genetics 206, 1699–1725 (2017).

2. Schuettengruber, B., Bourbon, H.-M., Croce, L. D. & Cavalli, G. Genome Regulation by Polycomb and Trithorax: 70 Years and Counting. Cell 171, 34–57 (2017).

3. Béguelin, W. et al. EZH2 is required for germinal center formation and somatic EZH2 mutations promote lymphoid transformation. Cancer Cell 23, 677–692 (2013).

4. Béguelin, W. et al. EZH2 and BCL6 Cooperate to Assemble CBX8-BCOR Complex to Repress Bivalent Promoters, Mediate Germinal Center Formation and Lymphomagenesis. Cancer Cell 30, 197–213 (2016).

5. Chan, H. L. et al. Polycomb complexes associate with enhancers and promote oncogenic transcriptional programs in cancer through multiple mechanisms. Nat Commun 9, 3377 (2018).

6. Donaldson-Collier, M. C. et al. EZH2 oncogenic mutations drive epigenetic, transcriptional, and structural changes within chromatin domains. Nat Genet 51, 517–528 (2019).

7. Duan, R., Du, W. & Guo, W. EZH2: a novel target for cancer treatment. J Hematol Oncol 13, 104 (2020).

8. Piunti, A. & Shilatifard, A. The roles of Polycomb repressive complexes in mammalian development and cancer. Nat Rev Mol Cell Biol 22, 326–345 (2021).

9. Schlesinger, Y. et al. Polycomb-mediated methylation on Lys27 of histone H3 pre-marks genes for de novo methylation in cancer. Nat Genet 39, 232–236 (2007).

10. Yap, D. B. et al. Somatic mutations at EZH2 Y641 act dominantly through a mechanism of selectively altered PRC2 catalytic activity, to increase H3K27 trimethylation. Blood 117, 2451–2459 (2011).

11. Cao, R. et al. Role of histone H3 lysine 27 methylation in Polycomb-group silencing. Science 298, 1039–1043 (2002).

12. Eskeland, R. et al. Ring1B Compacts Chromatin Structure and Represses Gene Expression Independent of Histone Ubiquitination. Mol Cell 38, 452–464 (2010).

13. Gao, Z., et al. PCGF homologs, CBX proteins, and RYBP define functionally distinct PRC1 family complexes. Mol Cell 45, 344–356 (2012).

14. Wang, H. et al. Role of histone H2A ubiquitination in Polycomb silencing. Nature 431, 873– 878 (2004).

15. Bernstein, B. E. et al. A bivalent chromatin structure marks key developmental genes in embryonic stem cells. Cell 125, 315–326 (2006).

16. Wang, L. et al. Hierarchical Recruitment of Polycomb Group Silencing Complexes. Molecular Cell 14, 637–646 (2004).

17. Blackledge, N. P. & Klose, R. J. The molecular principles of gene regulation by Polycomb repressive complexes. Nat Rev Mol Cell Biol 1–19 (2021) doi:10.1038/s41580-021-00398-y.

18. Blackledge, N. P. et al. PRC1 Catalytic Activity Is Central to Polycomb System Function. Molecular Cell 77, 857–874.e9 (2020).

19. Fursova, N. A. et al. Synergy between Variant PRC1 Complexes Defines Polycomb-Mediated Gene Repression. Mol Cell 74, 1020–1036.e8 (2019).

20. Kasinath, V. et al. JARID2 and AEBP2 regulate PRC2 in the presence of H2AK119ub1 and other histone modifications. Science 371, eabc3393 (2021).

21. Bantignies, F. et al. Polycomb-dependent regulatory contacts between distant Hox loci in Drosophila. Cell 144, 214–226 (2011).

22. Boyle, S. et al. A central role for canonical PRC1 in shaping the 3D nuclear landscape. Genes Dev. (2020) doi:10.1101/gad.336487.120.

23. Eagen, K. P., Aiden, E. L. & Kornberg, R. D. Polycomb-mediated chromatin loops revealed by a subkilobase-resolution chromatin interaction map. PNAS 114, 8764–8769 (2017).

24. Isono, K. et al. SAM domain polymerization links subnuclear clustering of PRC1 to gene silencing. Dev Cell 26, 565–577 (2013).

25. Kundu, S. et al. Polycomb Repressive Complex 1 Generates Discrete Compacted Domains that Change during Differentiation. Mol Cell 65, 432–446.e5 (2017).

26. Rhodes, J. D. P. et al. Cohesin Disrupts Polycomb-Dependent Chromosome Interactions in Embryonic Stem Cells. Cell Reports 30, 820–835.e10 (2020).

27. Schoenfelder, S. et al. Polycomb repressive complex PRC1 spatially constrains the mouse embryonic stem cell genome. Nat Genet 47, 1179–1186 (2015).

28. Ogiyama, Y., Schuettengruber, B., Papadopoulos, G. L., Chang, J.-M. & Cavalli, G. Polycomb-Dependent Chromatin Looping Contributes to Gene Silencing during Drosophila Development. Mol Cell 71, 73–88.e5 (2018).

29. Scelfo, A. et al. Functional Landscape of PCGF Proteins Reveals Both RING1A/B-Dependent- and RING1A/B-Independent-Specific Activities. Molecular Cell 74, 1037–1052.e7 (2019).

30. Zepeda-Martinez, J. A. et al. Parallel PRC2/cPRC1 and vPRC1 pathways silence lineage-specific genes and maintain self-renewal in mouse embryonic stem cells. Sci Adv 6, eaax5692 (2020).

31. Moussa, H. F. et al. Canonical PRC1 controls sequence-independent propagation of Polycomb-mediated gene silencing. Nat Commun 10, 1931 (2019).

32. Michlits, G. et al. CRISPR-UMI: single-cell lineage tracing of pooled CRISPR-Cas9 screens. Nat Methods 14, 1191–1197 (2017).

33. Li, W. et al. MAGeCK enables robust identification of essential genes from genome-scale CRISPR/Cas9 knockout screens. Genome Biol 15, 554 (2014).

34. Haering, C. H., Farcas, A.-M., Arumugam, P., Metson, J. & Nasmyth, K. The cohesin ring concatenates sister DNA molecules. Nature 454, 297–301 (2008).

35. Arruda, N. L. et al. Distinct and overlapping roles of STAG1 and STAG2 in cohesin localization and gene expression in embryonic stem cells. Epigenetics & Chromatin 13, 32 (2020).

36. Gandhi, R., Gillespie, P. J. & Hirano, T. Human Wapl is a cohesin-binding protein that promotes sister-chromatid resolution in mitotic prophase. Curr Biol 16, 2406–2417 (2006).

37. Haarhuis, J. H. I. et al. The Cohesin Release Factor WAPL Restricts Chromatin Loop Extension. Cell 169, 693–707.e14 (2017).

38. Huis in ‘t Veld, P. J., et al. Characterization of a DNA exit gate in the human cohesin ring. Science 346, 968–972 (2014).

39. Kueng, S. et al. Wapl controls the dynamic association of cohesin with chromatin. Cell 127, 955–967 (2006).

40. Losada, A., Yokochi, T., Kobayashi, R. & Hirano, T. Identification and characterization of SA/Scc3p subunits in the Xenopus and human cohesin complexes. J Cell Biol 150, 405–416 (2000).

41. Tedeschi, A. et al. Wapl is an essential regulator of chromatin structure and chromosome segregation. Nature 501, 564–568 (2013).

42. Wutz, G. et al. Topologically associating domains and chromatin loops depend on cohesin and are regulated by CTCF, WAPL, and PD S5 proteins. EMBO J 36, 3573–3599 (2017).

43. Zhang, N. et al. Characterization of the interaction between the cohesin subunits Rad21 and SA1/2. PLoS One 8, e69458 (2013).

44. Cuadrado, A. et al. Specific Contributions of Cohesin-SA1 and Cohesin-SA2 to TADs and Polycomb Domains in Embryonic Stem Cells. Cell Reports 27, 3500–3510.e4 (2019).

45. Elling, U. et al. A reversible haploid mouse embryonic stem cell biobank resource for functional genomics. Nature 550, 114–118 (2017).

46. Kagey, M. H. et al. Mediator and cohesin connect gene expression and chromatin architecture. Nature 467, 430–435 (2010).

47. Lavagnolli, T. et al. Initiation and maintenance of pluripotency gene expression in the absence of cohesin. Genes Dev. 29, 23–38 (2015).

48. Nitzsche, A. et al. RAD21 cooperates with pluripotency transcription factors in the maintenance of embryonic stem cell identity. PLoS One 6, e19470 (2011).

49. Chen, S., Jiao, L., Liu, X., Yang, X. & Liu, X. A Dimeric Structural Scaffold for PRC2-PCL Targeting to CpG Island Chromatin. Mol Cell 77, 1265–1278.e7 (2020).

50. Deaton, A. M. & Bird, A. CpG islands and the regulation of transcription. Genes Dev. 25, 1010–1022 (2011).

51. Farcas, A. M. et al. KDM2B links the Polycomb Repressive Complex 1 (PRC1) to recognition of CpG islands. Elife 1, e00205 (2012).

52. He, J. et al. Kdm2b maintains murine embryonic stem cell status by recruiting PRC1 complex to CpG islands of developmental genes. Nat Cell Biol 15, 373–384 (2013).

53. Li, H. et al. Polycomb-like proteins link the PRC2 complex to CpG islands. Nature 549, 287– 291 (2017).

54. Wu, X., Johansen, J. V. & Helin, K. Fbxl10/Kdm2b recruits polycomb repressive complex 1 to CpG islands and regulates H2A ubiquitylation. Mol Cell 49, 1134–1146 (2013).

55. Long, H. K. et al. Epigenetic conservation at gene regulatory elements revealed by non-methylated DNA profiling in seven vertebrates. eLife 2, e00348 (2013).

56. Busslinger, G. A. et al. Cohesin is positioned in mammalian genomes by transcription, CTCF and Wapl. Nature 544, 503–507 (2017).

57. Rao, S. S. P. et al. Cohesin Loss Eliminates All Loop Domains. Cell 171, 305–320.e24 (2017).

58. Knight, P. A. & Ruiz, D. A fast algorithm for matrix balancing. IMA Journal of Numerical Analysis 33, 1029–1047 (2013).

59. Bonev, B. et al. Multiscale 3D Genome Rewiring during Mouse Neural Development. Cell 171, 557–572.e24 (2017).

60. Crane, E. et al. Condensin-Driven Remodeling of X-Chromosome Topology during Dosage Compensation. Nature 523, 240–244 (2015).

61. Cunningham, M. D. et al. Wapl antagonizes cohesin binding and promotes Polycomb-group silencing in Drosophila. Development 139, 4172–4179 (2012).

62. Zhang, B. et al. Dosage Effects of Cohesin Regulatory Factor PDS5 on Mammalian Development: Implications for Cohesinopathies. PLoS One 4, e5232 (2009).

63. Katoh-Fukui, Y. et al. Male-to-female sex reversal in M33 mutant mice. Nature 393, 688– 692 (1998).

64. Lau, M. S. et al. Mutation of a nucleosome compaction region disrupts Polycomb-mediated axial patterning. Science 355, 1081–1084 (2017).

65. Cohen, I. et al. PRC1 preserves epidermal tissue integrity independently of PRC2. Genes Dev. 33, 55–60 (2019).

66. Pasini, D., Bracken, A. P., Hansen, J. B., Capillo, M. & Helin, K. The Polycomb Group Protein Suz12 Is Required for Embryonic Stem Cell Differentiation. Molecular and Cellular Biology 27, 3769–3779 (2007).

67. Riising, E. M. et al. Gene silencing triggers polycomb repressive complex 2 recruitment to CpG islands genome wide. Mol Cell 55, 347–360 (2014).

68. Akasaka, T. et al. Mice doubly deficient for the Polycomb Group genes Mel18 and Bmi1 reveal synergy and requirement for maintenance but not initiation of Hox gene expression. Development 128, 1587–1597 (2001).

69. Hansen, K. H. et al. A model for transmission of the H3K27me3 epigenetic mark. Nat Cell Biol 10, 1291–1300 (2008).

70. Chambeyron, S. & Bickmore, W. A. Chromatin decondensation and nuclear reorganization of the HoxB locus upon induction of transcription. Genes Dev 18, 1119–1130 (2004).

71. Illingworth, R. S. Chromatin folding and nuclear architecture: PRC1 function in 3D. Curr Opin Genet Dev 55, 82–90 (2019).

72. Adane, B. et al. STAG2 loss rewires oncogenic and developmental programs to promote metastasis in Ewing sarcoma. Cancer Cell 39, 827–844.e10 (2021).

73. Kubo, N. et al. Promoter-proximal CTCF binding promotes distal enhancer-dependent gene activation. Nat Struct Mol Biol 28, 152–161 (2021).

74. Nora, E. P. et al. Targeted Degradation of CTCF Decouples Local Insulation of Chromosome Domains from Genomic Compartmentalization. Cell 169, 930–944.e22 (2017).

75. Rowley, M. J. & Corces, V. G. Organizational principles of 3D genome architecture. Nat Rev Genet 19, 789–800 (2018).

76. Schwarzer, W. et al. Two independent modes of chromatin organization revealed by cohesin removal. Nature 551, 51–56 (2017).

77. Rao, S. S. P. et al. A 3D Map of the Human Genome at Kilobase Resolution Reveals Principles of Chromatin Looping. Cell 159, 1665–1680 (2014).

78. Dobin, A. et al. STAR: ultrafast universal RNA-seq aligner. Bioinformatics 29, 15–21 (2013).

79. Anders, S., Pyl, P. T. & Huber, W. HTSeq--a Python framework to work with high-throughput sequencing data. Bioinformatics 31, 166–169 (2015).

80. Love, M. I., Huber, W. & Anders, S. Moderated estimation of fold change and dispersion for RNA-seq data with DESeq2. Genome Biol 15, 550 (2014).

81. Zhu, A., Ibrahim, J. G. & Love, M. I. Heavy-tailed prior distributions for sequence count data: removing the noise and preserving large differences. Bioinformatics 35, 2084–2092 (2019).

82. Wu, T. et al. clusterProfiler 4.0: A universal enrichment tool for interpreting omics data. The Innovation 2, 100141 (2021).

83. Yu, G., Wang, L.-G., Han, Y. & He, Q.-Y. clusterProfiler: an R Package for Comparing Biological Themes Among Gene Clusters. OMICS: A Journal of Integrative Biology 16, 284– 287 (2012).

84. Li, H. et al. The Sequence Alignment/Map format and SAMtools. Bioinformatics 25, 2078– 2079 (2009).

85. Bonhoure, N. et al. Quantifying ChIP-seq data: a spiking method providing an internal reference for sample-to-sample normalization. Genome Res 24, 1157–1168 (2014).

86. Hu, B. et al. Biological chromodynamics: a general method for measuring protein occupancy across the genome by calibrating ChIP-seq. Nucleic Acids Res 43, e132 (2015).

87. Ramírez, F. et al. deepTools2: a next generation web server for deep-sequencing data analysis. Nucleic Acids Res 44, W160–W165 (2016).

88. Zhang, Y. et al. Model-based Analysis of ChIP-Seq (MACS). Genome Biology 9, R137 (2008).

89. Servant, N. et al. HiC-Pro: an optimized and flexible pipeline for Hi-C data processing. Genome Biology 16, 259 (2015).

90. Langmead, B. & Salzberg, S. L. Fast gapped-read alignment with Bowtie 2. Nat Methods 9, 357–359 (2012).

91. Ramírez, F. et al. High-resolution TADs reveal DNA sequences underlying genome organization in flies. Nat Commun 9, 189 (2018).

92. Wolff, J. et al. Galaxy HiCExplorer: a web server for reproducible Hi-C data analysis, quality control and visualization. Nucleic Acids Research 46, W11–W16 (2018).

93. Wolff, J. et al. Galaxy HiCExplorer 3: a web server for reproducible Hi-C, capture Hi-C and single-cell Hi-C data analysis, quality control and visualization. Nucleic Acids Research 48, W177–W184 (2020).

94. Durand, N. C. et al. Juicer Provides a One-Click System for Analyzing Loop-Resolution Hi-C Experiments. cels 3, 95–98 (2016).

95. Flyamer, I. M., Illingworth, R. S. & Bickmore, W. A. Coolpup.py: versatile pile-up analysis of Hi-C data. Bioinformatics 36, 2980–2985 (2020).

96. Weide, R. H. van der et al. Hi-C Analyses with GENOVA: a case study with cohesin variants. 2021.01.22.427620 https://www.biorxiv.org/content/10.1101/2021.01.22.427620v1 2021) doi:10.1101/2021.01.22.427620.

97. Kaaij, L. J. T., Mohn, F., van der Weide, R. H., de Wit, E. & Bühler, M. The ChAHP Complex Counteracts Chromatin Looping at CTCF Sites that Emerged from SINE Expansions in Mouse. Cell 178, 1437–1451.e14 (2019).

